# Stable neutralization of virulent bacteria using temperate phage in the mammalian gut

**DOI:** 10.1101/794222

**Authors:** Bryan B. Hsu, Jeffrey C. Way, Pamela A. Silver

## Abstract

Elimination or alteration of select members of the gut microbiota is key to therapeutic efficacy. However, the complexity of these microbial inhabitants makes it challenging to precisely target bacteria without unexpected cascading effects. Here, we use bacteriophage to deliver exogenous genes to specific bacteria by genomic integration of temperate phage for long-lasting modification. As a real-world therapeutic test, we engineered λ phage to transcriptionally-repress shigatoxin by using genetic hybrids between λ and other lambdoid phages to overcome resistance encoded by the virulent prophage derived from enterohemorrhagic *E. coli*. We show that a single dose of engineered phage propagates throughout the bacterial community and reduces shigatoxin production in an enteric mouse model of infection without markedly affecting bacterial concentrations. We thus minimize the selection for resistance by relying on anti-virulence and not anti-bacterial action. Our work reveals a new framework for transferring functions to bacteria within their native environment.

## INTRODUCTION

The human gut microbiota is a collection of microbes colonizing the gastrointestinal tract and has been associated with various aspects of human health.^1^ While this community typically works in concert with our bodies, substantial perturbations such as antibiotics or infections can disrupt the microbial balance and lead to long lasting dysbiosis.^2^ In some instances pathogenic bacteria do so by transmitting virulence factors encoded by these pathogens to commensal bacteria through plasmid-based^3^ and phage-based horizontal gene transfer (HGT).^4^ Remediating diseases associated with these pathogens while minimizing unintended and disruptive effects to the surrounding microbiota remains challenging^5^ especially with the limited tools available for targeting particular species.^6^

Our ability to manipulate the composition and function of the gut microbiota is presently limited in terms of precision and durability.^6^ Antibiotics non-specifically decimate swaths of gut species,^7^ dietary changes affect both the overall microbiota and mammalian host, probiotics poorly engraft due to colonization resistance,^8^ and even highly-specific lytic phages can cause cascading effects in the bacterial community despite targeting specific species.^9^ While in some cases these strategies may show transient efficacy, the emergence of resistant mutants can impact therapeutic effect. Broadly resetting the gut microbiota through fecal microbiota transplants (FMTs) has been promising especially for treating *Clostridium difficile* infections,^10^ but they are difficult to characterize and may transmit unintended traits such as obesity.^11^

An alternative strategy is to modify bacterial function within its native environment. For example, one approach has been to develop drugs that target the virulence factors of anti-biotic resistant pathogens to specifically neutralize their deleterious effects and minimizing selection for resistance. While a number of anti-virulence drugs are under investigation the targets for inhibition are generally limited to those accessible by small molecules and biologics (i.e., surface-bound and secreted proteins), may require multiple drugs targeting multiple virulence factors, and could have off target effects on other microbes and the host.^12^ While the principle of anti-virulence is attractive, it remains challenging in application.

Shigatoxin (Stx)-producing *E. coli* is one example of a pathogenic infection that is challenging to treat. Anti-virulence drugs targeting the toxin have been investigated but failed clinical trial.^13^ Antibiotics are contraindicated because of their potential to exacerbate virulence.^14^ While there are multiple virulence factors in the foodborne pathogen enterohemorrhagic *E. coli* (EHEC), Stx is significantly associated with disease severity^15^ and can lead to hemolytic uremic syndrome.^16^ Of the two main Stx variants—Stx1 and Stx2—the latter is ∼1000-fold more toxic.^17^ Similar to a number of other prophage-encoded virulence factors,^18^ Stx is not expressed while the phage is in a lysogenic state, *i.e.* stably integrated into the bacterial genome. It is not until induction, whether occurring spontaneously or from stimuli such as antibiotics, that the lytic life cycle is activated to produce Stx2^19^ and progeny phage that can spread virulence genes to commensal *E. coli* species.^20^

Instead of an antimicrobial strategy for killing pathogens, a genetic-based anti-virulence strategy could neutralize virulence before expression and minimize resistance until the bacteria has been completely shed from the gastrointestinal tract. Temperate phages offer a solution as they are genetically engineerable and can integrate into the bacterial chromosome as prophages for long-lasting effect as they confer fitness advantages to the bacterial host.^21^ Instead of relying on a non-native constituent of the gut that could face practical barriers for efficacy, temperate phages are abundantly found in human gut bacteria^22–24^ and can constitute large portions of the bacterial chromosome.^25^

To modify the function of specific species within the complex community in the mammalian gut, in a manner that avoids the emergence of resistance, we report the use of a genetically-engineered temperate phage to repress Stx from an established *E. coli* infection. We first show that genetic hybrids between lambdoid phages can overcome phage resistance mechanisms while maintaining function. We then genetically encode a transcriptional repressor of Stx in our engineered phage and show that it substantially reduces Stx produced by *E. coli in vitro*. Finally, we demonstrate that our engineered phage, when administered to mice pre-colonized by Stx-producing *E. coli*, can propagate throughout the murine gut from a single dose to significantly reduce fecal Stx concentrations. Our work describes a new therapeutic framework for the *in situ* modification of gut bacteria for genetic-based anti-virulence.

## RESULTS

### Temperate phage lysogenizes gut bacteria

Phages typically replicate by a lytic mechanism in which a bacterium infected by a phage turns its cellular machinery towards producing phage components that assemble into viral particles and are released upon cell lysis. This produces hundreds of progeny phage to infect new bacterial hosts (Figure A, *i-iii*). Lytic phages replicate solely by this life cycle and the decimation of phage-susceptible bacteria selects for phage-resistant mutants that can repopulate over time (Figure 1A, *upper panel*). In contrast, temperate phages can also integrate their genetic material into the host chromosome as a prophage to co-replicate with the bacterial genome during cell proliferation (Figure 1A, *iv-v*). In a bacterial population, this leads to the lysogenic conversion of phage-susceptible species that coexist with phage-resistant species (Figure 1A, *lower panel*). Because anti-bacterial approaches can enrich for resistance, including phage therapy which typically utilizes lytic phages, we aimed to engineer temperate phages to deliver an anti-virulence payload that neutralizes virulence in a manner that minimizes the selection for resistance.

**Figure 1.**
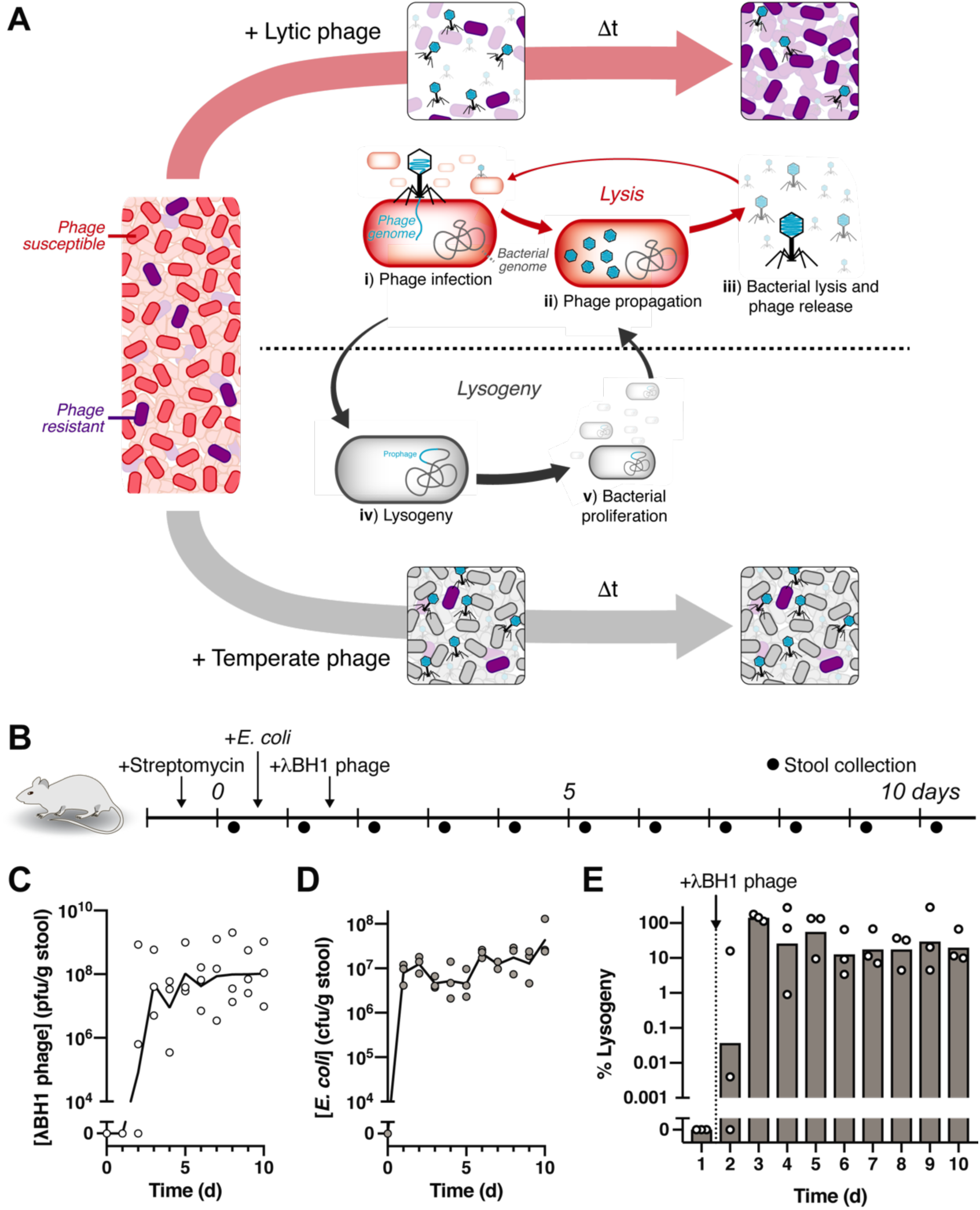
Temperate phage robustly transduces *E. coli* colonizing the mouse gut. (A, *upper panel*) Lytic phages decimate a bacterial population, selecting for resistant mutants that can repopulate over time. i) In lytic phage infection, the phage particle injects its genetic material into a bacterium and ii) directs the cell to produce phage components, iii) which are released upon cell lysis to continue infection. (A, *lower panel*) Temperate phages can infect a bacterial population to lysogenize phage-susceptible bacteria which persists over time. iv) Temperate phages can also participate in the lysogenic life cycle where they integrate their DNA into the bacterial genome and v) remain as a prophage during bacterial proliferation with the possibility of entering lytic replication in the future. (B) Experimental timeline examining the impact of λBH1 phage on pre-colonized *E. coli* in mice. (C) Fecal concentrations of free λBH1 phage and (D) *E. coli*. (E) Percentage of fecal *E. coli* lysogenized by λBH1. Symbols represent distinct samples from individual mice (*n* = 3) with lines or bars representing the geometric mean.

To illustrate the feasibility of using a temperate phage, we show that bacteriophage λ transduces a substantial fraction of targeted bacteria in the mammalian gut. As shown in Figure 1B, we used a streptomycin-treated mouse model to quantitate temperate phage lysogeny on *E. coli* colonizing the mammalian gastrointestinal tract. One day after colonization with *E. coli* MG1655, we introduced λBH1 phage by oral gavage and collected daily stool samples for analysis of bacterial and phage titers. We constructed λBH1 from λ phage by inserting an antibiotic resistance cassette for quantification of lysogens (Figure 2D). After oral administration of λBH1 phage, we found that fecal phage levels reached equilibrium approximately two days later and persisted at substantial concentrations (> 10^6^ pfu/g stool) for the duration of the experiment (Figure 1C). As phage in the absence of its cognate bacterial host is undetectable in the stool of mice ∼2 days after administration,^26^ our results indicate that λBH1 phage is capable of continuous replication in the gut, enabling its expansion throughout the bacterial population from a single dose. Furthermore, introduction of λBH1 phage did not significantly alter fecal *E. coli* concentrations (Figure 1D), which is in sharp contrast to lytic phages that can cause substantial reduction.^9^ Using antibiotic selection, we quantified the number of fecal *E. coli* harboring the λBH1 prophage and found a substantial fraction (∼17 to 30%) remained lysogenized by days 7 to 10 (Figure 1E). Overall, these results indicate that the temperate phage λ is capable of widespread modification of its cognate bacteria in the gut.

**Figure 2.**
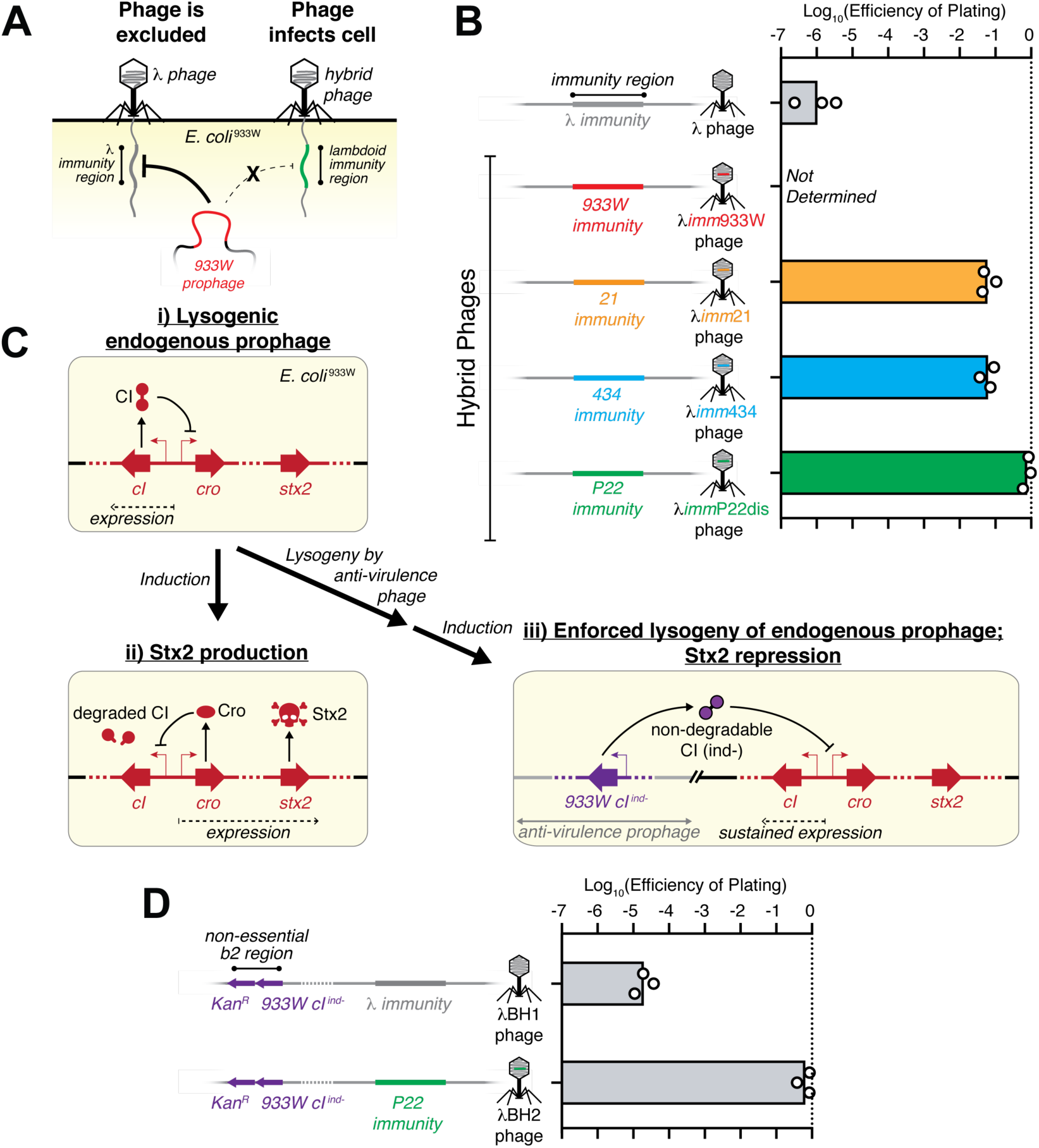
Hybrid λ phages overcome superinfection exclusion by prophage 933W. (A) Depiction of superinfection exclusion by the 933W prophage that inhibits infection by λ phage but is ineffective against a hybrid phage that contains the immunity region from another lambdoid phage in a λ phage background. (B) Schematic representation of a portion of the λ phage genome containing the λ immunity region and its hybrids containing immunity regions from other lambdoid phages (933W, 21, 434, and P22) in a λ phage background. Efficiency of plating (EOP) for λ phage and its hybrids on *E. coli* ^933W^ are shown to the right. (C) Genetic schemes of *E. coli* ^933W^ showing i) the 933W prophage expressing *cI* to maintain a lysogenic state in which *stx2* is not expressed, ii) induction that causes degradation of the CI protein leading to expression of the lytic genes including *cro* and *stx2*. This leads to cell lysis, releasing phage progeny and Stx2 protein. iii) Expression of a non-degradable cI for the 933W prophage, *933W.cI*^*ind-*^, from a genomically-integrated engineered temperate phage (anti-virulence prophage) can enforce the 933W prophage to remain lysogenic despite induction and degradation of endogenous CI protein. (D) Schematic representation of λBH1 phage, which is a λ phage with a kanamycin resistance cassette (*KanR*) and *933W.cI*^*ind-*^ inserted into the non-essential b2 region of the phage genome. λBH2 phage is a product of a phage cross between λBH1 and λ*imm*P22dis resulting in a phage containing *KanR* and *933W.cI*^*ind-*^ genes with a P22 immunity region in a λ phage background. Their respective EOP against *E. coli* ^933W^ are shown to the right. Symbols represent biological replicates with bars representing the geometric mean.

### Phage hybridization overcomes superinfection exclusion

As gut bacteria harbor numerous prophages including those encoding virulence,^27^ overcoming superinfection exclusion mechanisms is crucial for achieving efficacious *in situ* transduction. For the foodborne pathogen EHEC, the lambdoid prophage, 933W, both produces Stx2 and inhibits phage superinfection by other lambdoid phages.^28^ To demonstrate the efficacy of the *in situ* phage-based anti-virulence strategy, we aimed to neutralize Stx2 production from *E. coli* MG1655 lyosgenized by 933W (*E. coli* ^933W^).

*E. coli* ^933W^ excludes λ phage infection but genetic hybrids of λ with other lambdoid phages restore infectivity. As shown schematically in Figure 2A, the 933W prophage inhibits infection from λ phage by recognition of its immunity region, *i.e.* indispensable genes responsible for the lysis-lysogeny decision in the phage life cycle. Because lambdoid phages have similarities in genetic function and organization despite dissimilar sequences, it is feasible to replace the λ immunity region with orthologous immunity regions from other lambdoid phages to overcome the superinfection exclusion.^29^ We found that the efficiency of plating (EOP) of λ phage against *E. coli* ^933W^ was ∼10^6^-fold lower than that of the non-lysogen (Figure 2B), confirming its superinfection exclusion. We verified that this effect is not due to a cI-based immunity (Figure S1). We then tested the EOP for genetic hybrids of λ phage in which the λ immunity region is swapped with that of other lambdoid phages (*e.g.* 21, 434, and P22), (Figure S2 and Table S3). As shown in Figure 2B, these hybrid phages had substantially improved EOPs against *E. coli* ^933W^ with 6.0% for λ*imm*21 and 6.7% for λ*imm*434 phages. Moreover, hybridization with the Salmonella phage P22 resulted in near complete recovery of EOP to 78% for λ*imm*P22dis phage, indicating that lambdoid phages from non-cognate bacterial hosts could be a reservoir for genetic orthologs that maintain phage function while circumventing superinfection exclusion mechanisms.

### Genetic-based anti-virulence encoded by hybrid temperate phage

Expression of Stx2 is dependent upon induction of the 933W prophage. As schematically depicted in Figure 2C, *panel i)*, the 933W prophage in *E. coli* ^933W^ maintains a dormant state by expression of its repressor protein, *cl*, which blocks expression of *cro* and consequently lytic genes including Stx2. Induction, which occurs spontaneously and by stimuli such as antibiotics, causes activation of the bacterial SOS response and RecA-mediated degradation of Cl^30^ (Figure 2C, *panel ii*). This ultimately leads to expression of the lytic genes that produce phage progeny and Stx2. As the phage encoded repressor for the 933W prophage, *cl*, is key to blocking lytic induction and maintaining the dormant lysogenic state,^31^ constitutive expression of a non-degradable mutant of this repressor (*933W.cl*^*ind-*^) that contains a Lys178Asn mutation^31^ blocks induction of the 933W prophage and consequently neutralizes production of progeny phage and Stx2 (Figure 2C, *panel iii*), ultimately demonstrating anti-virulence at a genetic level.

Efficient gene transduction enables the delivery of anti-virulence genes. We inserted genes for *933W.cl*^*ind-*^ and a kanamycin resistant cassette (to quantitate lysogeny) into the non-essential b2 region of λ,^30^ producing λBH1 (Figures 2D and S3). We confirmed that *933W.cl*^*ind-*^ expressed from λBH1 was functional (Figure S4). To overcome superinfection exclusion from the 933W prophage, we utilized a P22 immunity region instead of a λ immunity region. A phage cross between λBH1 and λ*imm*P22dis resulted in the replacement of ∼6 kb of the immunity region of λBH1 with a ∼5 kb portion of that from λ*imm*P22dis while retaining *933W.cl*^*ind-*^ and *Kan*^*R*^ genes (Figures 2D and S3, and Table S3). This new phage, λBH2, showed improved EOP to 90% (Figure 2D) and demonstrated a functional loss of λ immunity and gain of P22 immunity, as well as expression of functional *933W.cl*^*ind-*^ (Figure S4).

### Anti-virulence phage inhibits Stx2 production *in vitro*

Transcriptional repression delivered by λBH2 phage neutralizes Stx2 production. As outlined in Figure 3A, we tested the efficacy of λBH2 phage to inhibit Stx2 production from *E. coli* ^933W^ by mixing them at equal concentrations (MOI∼1) and culturing for 8 h. We found significantly less Stx2 produced in *E. coli* ^933W^ cultures treated with λBH2 phage compared to those untreated (“buffer”) or treated with λ*imm*P22dis phage, the parental phage of λBH2 that is capable of infecting *E. coli* ^933W^ but lacks the *933W.cl*^*ind-*^ gene (Figure 3B). Quantification of bacterial concentrations over time show that *E. coli* ^933W^ steadily grows over 8 h in the absence of phage (“buffer”) whereas introduction of λ*imm*P22dis results in an initial drop in titer during the first 4 h followed by a recovery (Figure 4D, *non-induced*). For λBH2 phage, a similar drop in bacterial concentration was associated with increased lysogenic conversion of *E. coli* ^933W^ that reached 70% by 4 h, indicating that both decreased bacterial titers and repressed *Stx2* expression may contribute to the overall reduction of Stx2 concentration. To confirm that the latter provides sustained anti-virulence effect, we isolated λBH2 lysogens of *E. coli* ^933W^, *i.e. E. coli* containing prophages of both 933W and λBH2 (Figure 4F) and measured the Stx2 produced in culture. While an *E. coli* ^933W^ culture accumulated 13.1 ng/mL of Stx2 over 8 h, there were none detected for λBH2 lysogens (Figure 4G). Similarly, λBH1 lysogens do not produce detectable concentrations of Stx2 despite their poor ability to initially infect *E. coli* ^933W^, confirming that once lysogenic conversion occurs the resultant lysogens do not produce Stx2.

**Figure 3.**
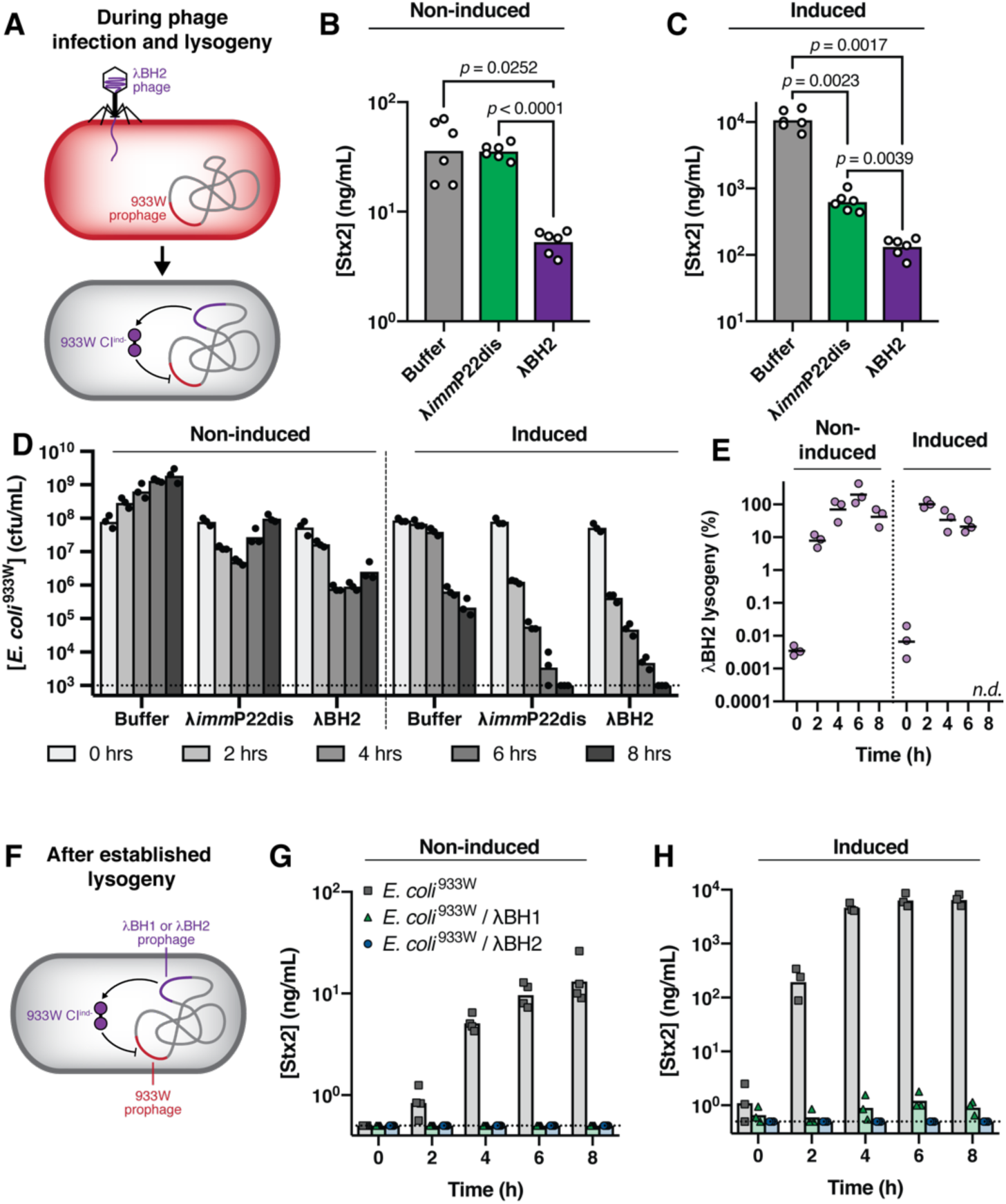
Engineered λ phage neutralizes shigatoxin production from *E. coli* ^933W^ *in vitro*. (A) *E. coli* ^933W^ was mixed with buffer, λ*imm*P22dis or λBH2 free phage (MOI∼1) at t = 0 from which (B) concentration of Stx2 was measured after 8 h of *in vitro* culture under non-induced and (C) induced conditions (0.5 μg/mL of mitomycin c). Significance was calculated by one-way ANOVA with post-hoc Tukey test. (D) Total *E. coli* ^933W^ and (E) the percentage of bacteria lysogenized by λBH2 was measured over 8 h under non-induced and mitomycin c induced conditions. (F) *E. coli* ^933W^ lysogenized with λBH1 or λBH2 was cultured *in vitro* and analyzed for Stx2 produced under (G) non-induced or (H) induced conditions. Symbols represent biological replicates with bars or lines representing the geometric mean.

**Figure 4.**
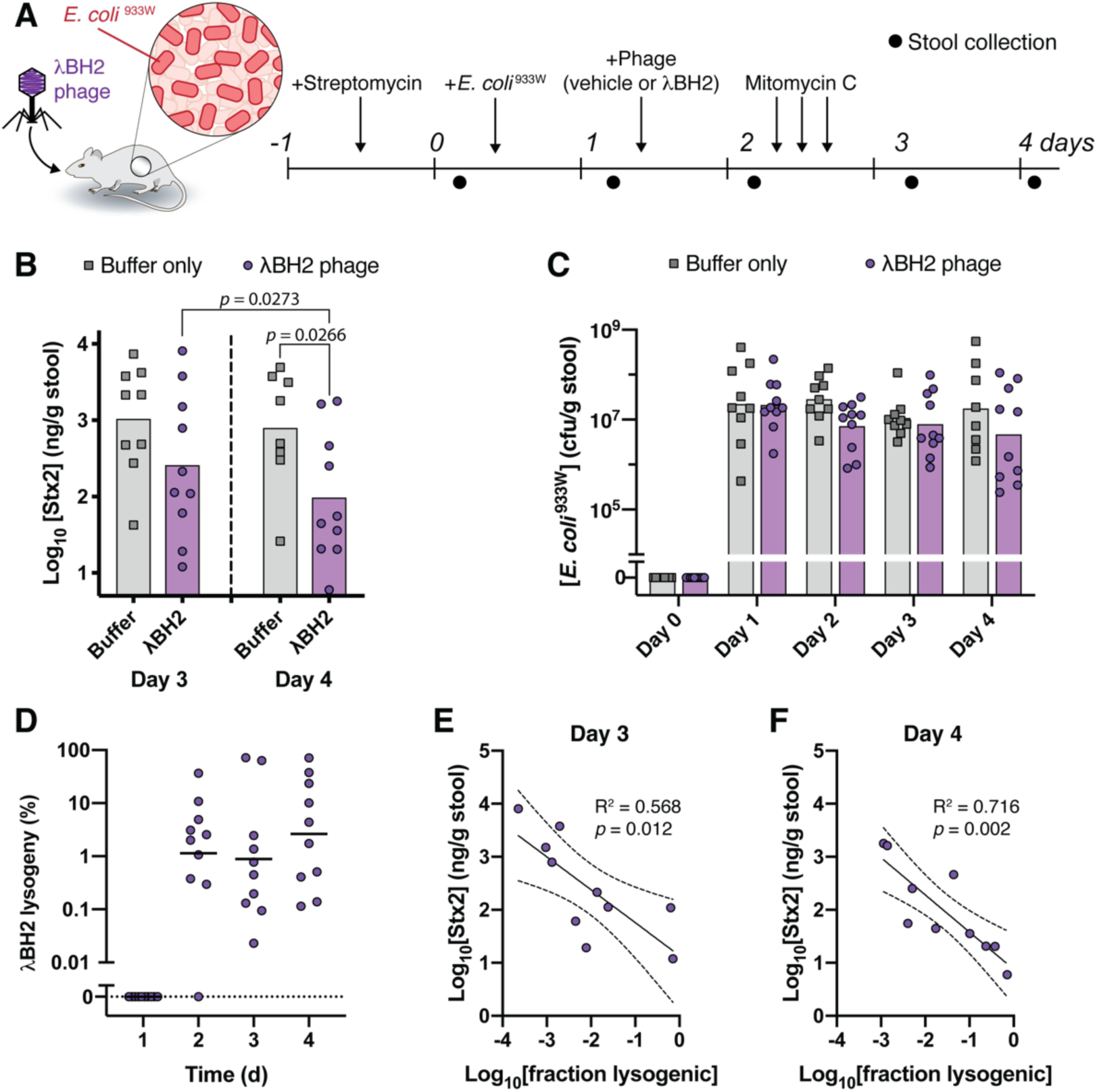
λBH2 phage lysogenizes *E. coli* ^933W^ in the murine gut and reduces fecal shigatoxin concentrations. (A) Streptomycin-treated mice pre-colonized with *E. coli* ^933W^ received one dose of 5 × 10^9^ pfu of λBH2 phage orally. Mitomycin c was administered thrice at 3 h intervals by intraperitoneal injection to induce Stx2 expression in the gut. (B) Concentrations of fecal Stx2 after induction with mitomycin c. Two-tailed Mann-Whitney tests were used to compare Buffer and λBH2 conditions while two-tailed Wilcoxon tests were used to compare between Day 3 and Day 4 within the same group. (C) Concentrations of total fecal *E. coli* ^933W^ and (D) percentage of fecal *E. coli* ^933W^ found to be lysogenized by λBH2 phage. (E) Concentration of fecal Stx2 as a function of fraction of fecal *E. coli* ^933W^ lysogenic for λBH2 phage on day 3 and (F) on day 4. Line and dashed lines represent mean and 95% confidence intervals of linear regression, respectively. *P* value describes significance of slope being non-zero. Symbols represent individual mice for Buffer (n = 9) and λBH2 conditions (n = 10). On Day 4, one buffer-treated mouse was unable to produce stool for analysis. Bars or lines represent geometric means.

Stx2 repression is maintained under inducing conditions. DNA damaging agents such as antibiotics can induce lambdoid prophages towards lysis (Figure 1E to 1B) by activating the bacterial SOS response leading to RecA-mediated degradation of CI.^31^ To test whether λBH2 phage remains effective under these more aggressive lytic conditions, we measured Stx2 produced in cultures of λBH2 phage mixed with *E. coli* ^933W^ (Figure 3A) in the presence of an inducing agent, mitomycin c. As shown in Figure 3C, *E. coli* ^933W^ receiving buffer alone produces substantially more Stx2 when incubated with mitomycin c due to induction of the 933W prophage. This induction also directs other phages towards primarily lytic replication and so the introduction of λ*imm*P22dis phage significantly reduces *E. coli* ^933W^ concentrations (Figure 3D, *induced*) and consequently Stx2 concentrations (Figure 3C). Ultimately, λBH2 phage treatment achieves significantly lower Stx2 concentrations than those measured for buffer and λ*imm*P22dis phage conditions (Figure 3C) because it is capable of repressing *stx2* expression from a large fraction of *E. coli* ^933W^ as shown by the substantial lysogenic conversion (Figure 3E, *induced*). To confirm that once lysogeny is established by λBH1 or λBH2 phage, *stx2* repression is maintained even in the presence of mitomycin c, we cultured λBH2 lysogens of *E. coli* ^933W^ for up to 8 h in the presence of mitomycin c (Figure 3F). In the case of λBH2 lysogens of *E. coli* ^933W^, we were unable to detect the toxin, indicating repression is maintained under inducing conditions (Figure 3H).

### Anti-virulence phage reduces Stx2 production *in vivo*

λBH2 reduces fecal Stx2 concentrations in mice. To determine if our phage-based anti-virulence strategy is effective *in vivo*, we used a mouse model of enteric Stx2 intoxication from Stx-producing *E. coli*.^32^ While it is challenging to model the effect of enteric pathogens including Stx-producing *E. coli* in mice,^33^ mitomycin c injections can induce substantial quantities of Stx that is otherwise too low to be detected in stool. Mice pre-colonized by *E. coli* ^933W^ were orally treated with λBH2 phage and then received three doses of mitomycin c by intraperitoneal injection to induce *stx2* expression (Figure 4A). Daily fecal samples were collected for analysis of bacterial and Stx2 concentrations. After mitomycin c injections, we quantified fecal Stx2 and found that λBH2 treatment reduced fecal Stx2 titers compared to buffer on days 3 and 4 with the latter showing statistical significance (Figure 4B). Furthermore, mice receiving λBH2 phage showed a significant reduction in fecal Stx2 from day 3 to 4, whereas buffer treated mice did not (*p* = 0.547; Wilcoxen test). Although λBH2 phage did not completely repress Stx2 production, these are highly inducing non-physiological conditions with fecal Stx2 concentrations (∼10^2^ to 10^3^ ng Stx2/g mouse stool) in excess of those encountered in human Stx-producing *E. coli* infections (∼2-50 ng Stx2/mL human stool).^1^

λBH2 phage lysogenizes *E. coli* ^933W^ and does not affect its titer in the murine gut. Quantification of total fecal *E. coli* ^933W^ did not reveal markedly different concentrations between buffer and λBH2 treated mice (Figure 4C). Quantification of fecal *E. coli* ^933W^ lysogenized by λBH2 (Figure 4D) showed that a substantial fraction of the population was transduced, with geometric means between ∼0.9% to 2.6% and individual samples reaching as high as 71%. Notably, there is a large spread in lysogeny between individual mice with daily means lower than what we found in our previous experiment with *E. coli* lacking the 933W prophage (Figure 1E). Past work has shown that a high level of induction disfavors lysogeny^35^ and within the context of this study, our use of a mitomycin c mouse model may underestimate the achievable degree of lysogeny in more typical, less-inducing conditions such as those shown in Figure 1B-1E.

Lysogeny by λBH2 phage reduces fecal concentrations of Stx2. Though treatment with λBH2 phage shows a reduction in the average fecal Stx2 between treated and untreated groups, we aimed to confirm whether this reduction is indeed associated with lysogenic conversion by λBH2 phage. By plotting the fecal concentration of Stx2 as a function of lysogeny by λBH2, we found that for both days 3 and 4 there is a significantly non-zero inverse correlation (Figures 4E and 4F) confirming that reduced fecal Stx2 is caused by anti-virulence effect from λBH2 phage.

## DISCUSSION

Here we demonstrate a genetic strategy for *in situ* anti-virulence treatment of bacteria colonizing the gut. We genetically engineer temperate phage λ to express a repressor that neutralizes Stx production in *E. coli* and take advantage of the genetic mosaicism of lambdoid phages to create a hybrid phage that is capable of overcoming phage resistance mechanisms. We found that our anti-virulence phage not only efficiently infects, lysogenizes and inhibits Stx2 production from *E. coli in vitro*, but is also effective at propagating in the murine gut from a single dose to significantly reduce Stx2 production *in vivo*.

With the complexity and interconnectedness of microbes in the gut, perturbations can have unexpected consequences. Modulating the impact of a bacterial species by manipulating its concentration can lead to unintended cascading effects mediated by inter-bacterial or bacterial-host interactions. While the typical strategy is to eliminate a particular bacteria, it is usually a specific function performed by this bacteria that is deleterious. By precisely and robustly modifying this individual function, a therapeutic effect can persist while minimizing disruption to the surrounding microbiota and avoiding the selection for resistance. Herein we report a framework for making precise genetic modifications that can be practically applied to bacteria within a complex biological system such as the gut microbiome.

For treating pathogenic bacterial infections, disarming their pathogenicity by targeting virulence factors provides a direct therapeutic strategy. With the impending crisis of antimicrobial resistant infections, new strategies for combating pathogens are desperately needed. By aiming to repress virulence factors instead of relying on anti-bacterial effect, the selection pressures for resistance are minimized. Furthermore, using a genetic-based approach makes it feasible to rationally design anti-virulence strategies that target one or multiple virulence factors to improve therapeutic efficacy.

As we gain greater insight into the importance of the gut microbiota to health and its inter-individual diversity and complexity, modification of specific bacteria within this community requires a thoughtful and nuanced approach that maximizes therapeutic efficacy and minimizes collateral effect. Our work illustrates a framework by which bacteria can be specifically modulated *in situ* with rationally designed function in a manner alternative to present approaches and could inspire strategies for treating recalcitrant bacterial infections.

## ACKNOWLEDGEMENTS

This project was funded by the Bill & Melinda Gates Foundation through the Grand Challenges Explorations Initiative (OPP1150555) and the Defense Advanced Research Program Agency (DARPA BRICS HR0011-15-C-0094). B.B.H. received support from the Rosenbloom postdoctoral fellowship. We are grateful to James Kuo and Elizabeth Libby for their critical reading of this manuscript.

## AUTHOR CONTRIBUTIONS

B.B.H. and J.C.W. designed experiments and analyzed data. B.B.H. performed the experiments. J.C.W. and P.A.S. supervised the research. B.B.H. and P.A.S. wrote the manuscript.

## MATERIALS AND METHODS

### Animal studies

Animal work was approved by the Harvard Medical School IACUC under protocol number 4966. Female BALB/c mice (Charles River Laboratories) 6-7 weeks old were acclimated for one week prior to experiments.

To study of the effect of temperate phage on non-pathogenic *E. coli* in the mouse gut (Figure 2), mice received 5 g/L of streptomycin sulfate (Gold Bio) in their drinking water which was replaced every 2-3 days. On day 0, 100 μL of streptomycin-resistant *E. coli* MG1655 was administered to mice by oral gavage. The bacterial gavage solution was prepared from an overnight culture in LB, washed twice with PBS, and then diluted 100-fold into PBS, yielding ∼10^7^ cfu/mL. One day later (day 1), mice received 100 μL of λBH1 phage which consisted of a 5 × 10^7^ pfu/mL solution diluted 1:10 into 100 mM sodium bicarbonate immediately prior to gavage. Daily stool samples were collected for microbial quantification. To quantify fecal phage, fresh non-frozen samples were gently suspended into 1 mL of phage buffer, incubated at 4°C for ∼10 min with a few drops of chloroform, and then pelleted at 4000 rpm at 4°C. Phage concentration was determined using a double-agar overlay plaque assay^36^ in which serially diluted phage solutions were incubated for 20 min at r.t. with a hardened overlay of *E. coli* MG1655 in 0.3% agar in TNT media over a 1.5% agar in TNT base. After aspiration, plates were incubated at 37°C overnight after which plaques were counted. To quantify fecal *E. coli*, frozen stool was thawed from −80°C and suspended into 1 mL of PBS by vortexing for 10 min at 4°C followed by low-speed centrifugation at 200 rpm for 20 min to settle fecal debris. The fecal suspension was then serially diluted into PBS and 100 μL was plated onto MacConkey agar (Remel) plates supplemented with 100 μg/mL streptomycin sulfate to quantify total *E. coli* or supplemented with 100 μg/mL streptomycin and 50 μg/mL kanamycin to quantify λBH1 lysogens of *E. coli*.

To study the effect of the engineered temperate phage on Stx2-producing *E. coli*, mice were treated with similar conditions as described above with the following modifications. On day 0, mice received 100 μL of similarly prepared streptomycin-resistant *E. coli* ^933W^ in PBS by oral gavage. On day 1, mice received 100 μL of λBH2 phage, which was a 5 × 10^10^ pfu/mL solution diluted 1:10 into 100 mM sodium bicarbonate immediately prior to gavage. On day 2, to induce Stx2 expression from engrafted *E. coli*, mice received three intraperitoneal injections of 0.25 mg/kg of mitomycin c at 3 h intervals.^32^ Stool samples were collected daily and stored at −80°C until analysis. Fecal *E. coli* was quantified by plating as described above and fecal Stx2 was quantified from the same suspension of stool in PBS by mixing 10:1 with 20 mg/mL of polymyxin B, incubating at 37°C for ∼20 min and then storing at −20°C until analysis by ELISA as described below.

### Bacterial strains

A table of bacteria used in this study is listed in Table S1. *E. coli* ^933W^ was generated by a previously described method,^37^ in which 933W phage was produced from the supernatant of a log phase culture of *E. coli* O157:H7 strain edl933 in a modified LB media (10 g/L tryptone, 5 g/L yeast extract, 5 mM sodium chloride, 10 mM calcium chloride, and 0.001% thiamine) and then stored at 4°C. Molten top agar containing 100 μL of *E. coli* MG1655 and 3 mL of modified LB media with 0.3% agar at 45°C was poured onto plates of modified LB agar and allowed to harden. Supernatants of *E. coli* O157:H7 cultures were then spotted onto the top agar and incubated at 37°C overnight. Resulting plaques were picked and re-streaked onto LB. Successful 933W lysogens of *E. coli* were identified by screens for resistance to λ*imm*933W and the PCR amplification of the *cI* to *cro* region of 933W (fwd-agccactcccttgcctcg; rev-gcttatttcaagcatttcgcttgc). *E. coli* lysogens of λ and λ*imm*933W were generated similarly using TNT media instead of modified LB media and screened for successful lysogeny by resistance to λ or λ*imm*933W, respectively, and ability to produce phage progeny.

### Preparation of high titer phage stocks

Phage was propagated via the double agar overlay method where 100 μL of serially-diluted phage in phage buffer was mixed with 100 μL of *E. coli* MG1655 for ∼20 min at r.t., then mixed with 3 mL of molten top agar (TNT media with 0.3% top agar at 45°C) and poured onto pre-warmed plates of TNT agar. After incubation overnight at 37°C, top agar from plates with the highest density of plaques were suspended into 5 mL of phage buffer and then gently rocked at 4°C for ∼2 hrs. Supernatants were sterile filtered to yield ∼10^9^ to 10^10^ pfu/mL of phage. Phage stocks were stored at 4°C.

### Phage strains

A table of phage used in this study is listed in Table S2. λBH1 phage was generated using the inherent λred recombination system of λ phage expressed during lytic replication. A crude phage lysate containing recombinant phage was produced by mixing 100 μL of *E. coli* C600, containing a plasmid vector with Tn5-933W.cl^ind-^ flanked by 400 bp homology to ea59 and ea47 in a pJET1.2 backbone (Table S3), with 100 μL of serially-diluted λ phage, incubated for 20 min at r.t. followed by addition with 3 mL of molten top agar (TNT media with 0.3% top agar at 45°C) and poured onto TNT agar plates. After overnight incubation at 37°C, top agar from the plate with the greatest plaque density was resuspended into 5 mL of phage buffer (50 mM tris, 100 mM sodium chloride, 10 mM magnesium sulfate, and 0.01% gelatin, pH 7.5), sterile filtered, and stored at 4°C. To isolate the recombinant phage, 50 μL of crude phage lysate was mixed with 50 μL of *E. coli* C600 grown to log-phase in LB and incubated for ∼3 h at 37°C. After incubation, 100 μL was plated onto LB containing 50 μg/mL of kanamycin and grown overnight at 37°C with individual colonies re-streaked twice. To additionally purify by plaque purification, colonies were grown overnight in LB and their sterile filtered supernatants were spotted onto TNT top agar of *E. coli* C600. Individual plaques were streaked onto LB containing 50 μg/mL of kanamycin and sequenced to confirm insertion in the correct locus of λ phage.

λBH2 phage was generated by a phage cross between λBH1 phage and λ*imm*P22dis phage. 200 μL of a log phase *E. coli* C600 culture (7 × 10^7^ cfu/mL) in T-broth (1% tryptone and 0.5% sodium chloride) with 0.4% maltose was mixed with a 200 μL solution of λBH1 phage (1.5 × 10^8^ pfu/mL) and λ*imm*P22dis (1.5 × 10^8^ pfu/mL) in phage buffer. After static incubation at 37°C for 20 min, this mixture was diluted 100-fold into pre-warmed T-broth with 1% glucose and cultured with shaking at 37°C for 90 min. The culture was treated with drops of chloroform, pelleted, and then the supernatant was sterile-filtered to produce a crude phage lysate. Residual chloroform was minimized by crossflowing air (Millipore Steriflip) at r.t. for 1 h. 10 mL of this phage lysate was mixed with 1 mL of mid-log culture of a λ lysogen of *E. coli* C600 grown in LB with 0.4% maltose, incubated at 37°C for 20 min. After ∼30-fold concentration by centrifugation, 200 μL was plated onto LB containing 50 μg/mL of kanamycin. Resulting colonies were re-streaked twice and then tested for phage immunity by spot testing 5 μL of phage against a top agar containing each candidate colony. Correctly engineered phages, as lysogens, were identified from colonies by susceptibility to λ*imm*434 (positive control) but resistance to λ*imm*933W (presence of *933WcI*^*ind-*^ gene) and resistance to λ*imm*P22dis (presence of P22 immunity region). Phage lysates were prepared by culturing colonies overnight in TNT media, pelleting and sterile filtering the supernatant, and then incubating 100 μL of this phage mixture with 100 μL of *E. coli* C600 (MOI∼0.1) at 37°C for 20 min and then plating onto LB with 50 μg/mL of kanamycin. After overnight incubation at 37°C, phage was plaque purified by preparing phage lysates from individual colonies as described above and streaking 10 μL onto hardened top agar containing *E. coli* C600. After overnight incubation at 37°C, individual plaques were picked and restreaked onto LB with kanamycin. The resultant λBH2 lysogen of *E. coli* was confirmed susceptible to λ and λ*imm*434 as well as resistant to λ*imm*933W and λ*imm*P22dis (Figure S4). Sequencing confirmed the presence of the P22 immunity region and *933Wcl*^*ind-*^ gene (Figure S3 and Table S3).

### Quantifying phage and efficiency of plating

The infectivity of phage against *E. coli* was quantified by the double overlay agar method in which *E. coli* MG1655 or *E. coli* ^933W^ was cultured overnight in TNT media, diluted 1:100 into fresh TNT media and cultured until mid-log phase of which 50 μL was mixed with 700 μL of molten top agar (TNT media with 0.5% agar at 45°C) and poured into individual wells of a 6-well plate containing pre-poured TNT media with 1.5% agar. After hardening, 100 μL of phage serially-diluted in phage buffer was added and incubated for 20 min at r.t. followed by aspiration. Plates were incubated at 37°C overnight and then examined for titers of plaque forming units. Efficiency of plating was calculated as the titer of phage on the *E. coli* ^933W^ divided by its titer on the non-lysogenic *E. coli*.

### In vitro assay of phage effect

*E. coli* ^933W^ was cultured overnight in TNT media at 37°C, then cells were washed once with fresh TNT media and diluted to OD_600nm_ = 0.1 (∼8 × 10^7^ cfu/mL). At t = 0, 5 mL of *E. coli* suspension was mixed with 1 mL of 4 × 10^8^ pfu/mL of λBH2 or λ*imm*P22dis phage solution. To quantify *E. coli* concentrations in solution, aliquots were collected, serially-diluted into PBS and the spotted (10 μL) onto LB or LB with 50 μg/mL kanamycin plates to quantify total *E. coli* and λBH2 lysogens, respectively. After 8 h, aliquots were mixed 10:1 with 20 mg/mL of polymyxin B, incubated at 37°C for ∼20 min and stored at - 20°C for quantification of Stx2 by ELISA.

Stx2 concentrations in cultures of *E. coli* ^933W^, its λBH1 lysogen, or its λBH2 lysogen were prepared similarly as described to above, where overnight cultures were washed with fresh TNT media and diluted to OD_600nm_ = 0.1. During incubation at 37°C, aliquots were collected, mixed with polymyxin B and stored at −20°C for quantification of Stx2 by ELISA.

### ELISA quantification of Stx2

Maxisorp plates (ThermoScientific) were incubated with 100 μL/well of mouse monoclonal Stx2 antibody (Santa Cruz Biotechnology, sc-52727) diluted 1:2500 into PBS for 1.5 h at r.t.. Plates were thrice washed with PBST (PBS with 0.05% Tween20), then incubated with 200 μL of 1% BSA in PBS overnight at 4°C. After washing thrice with PBST, 100 μL/well of samples and a standard curve of diluted Stx2 (List Biological Labs, #164) were incubated at r.t. for 2 h. Following sample incubation, plates were washed thrice with PBST and incubation with 100 μL/well of anti-Stx2 Ab-HRP conjugate for 1 h at r.t.. The antibody-enzyme conjugate was previously prepared using an HRP conjugation kit (Abcam #ab102890) with a rabbit anti-Stx2 antibody (List Biological Labs, #765L) according to the manufacturer’s protocol. After washing thrice with PBST, 100 μL/well of colorimetric reagent Ultra TMB (ThermoFisher) was incubated at 37°C for 30 min prior to the addition of 50 μL/well of 2M H_2_SO_4_ to stop the reaction. Absorbance was measured at 450 nm.

## Data availability

All materials are readily available from the corresponding authors upon request or are commercially available. There are no restrictions on availability of the material used in the study.

