## Supplemental Information for "Stable neutralization of virulent bacteria using temperate phage in the mammalian gut"

**Figure S1**. Spot testing of λ and λ*imm*933W phage against lawns of their lysogens in *E. coli*. The zone of lysis (dark circle) on a lawn of E. coli indicates successful phage infection. Both phages are capable of infecting non-lysogens and lysogens of the other phage (i.e. λ phage infecting a λ*imm*933W lysogen and *vise-versa*) indicating the 933W·cI and λ·cI are not inhibitory to λ phage and 933W phage infection, respectively.

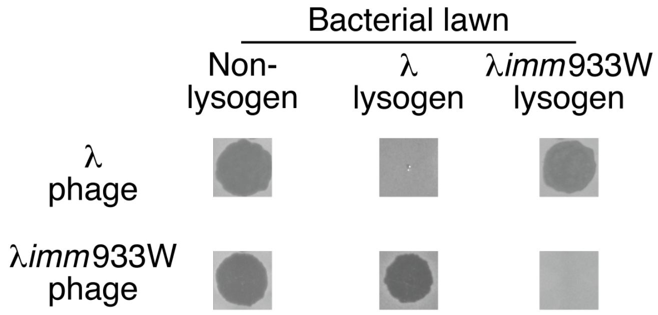

**Figure S2**. Genetic map of immunity regions within λ phage and its hybrids. (A) Genetic map of portions of the λ phage genome with brackets indicating the regions within λ that have been replaced with immunity regions from other lambdoid phages. (B) Within λ*imm*933W phage, the genes homologous to 933W (Accession ID NC.000924) are labeled in red and those homologous to λ (NC.001416) are labeled in black. Percent identities for genes less then identical are indicated in parenthesis. (C) Within λ*imm*P22dis phage, the genes homologous to P22 (AF217253) are labeled in green and those homologous to λ (NC.001416) are labeled in black.

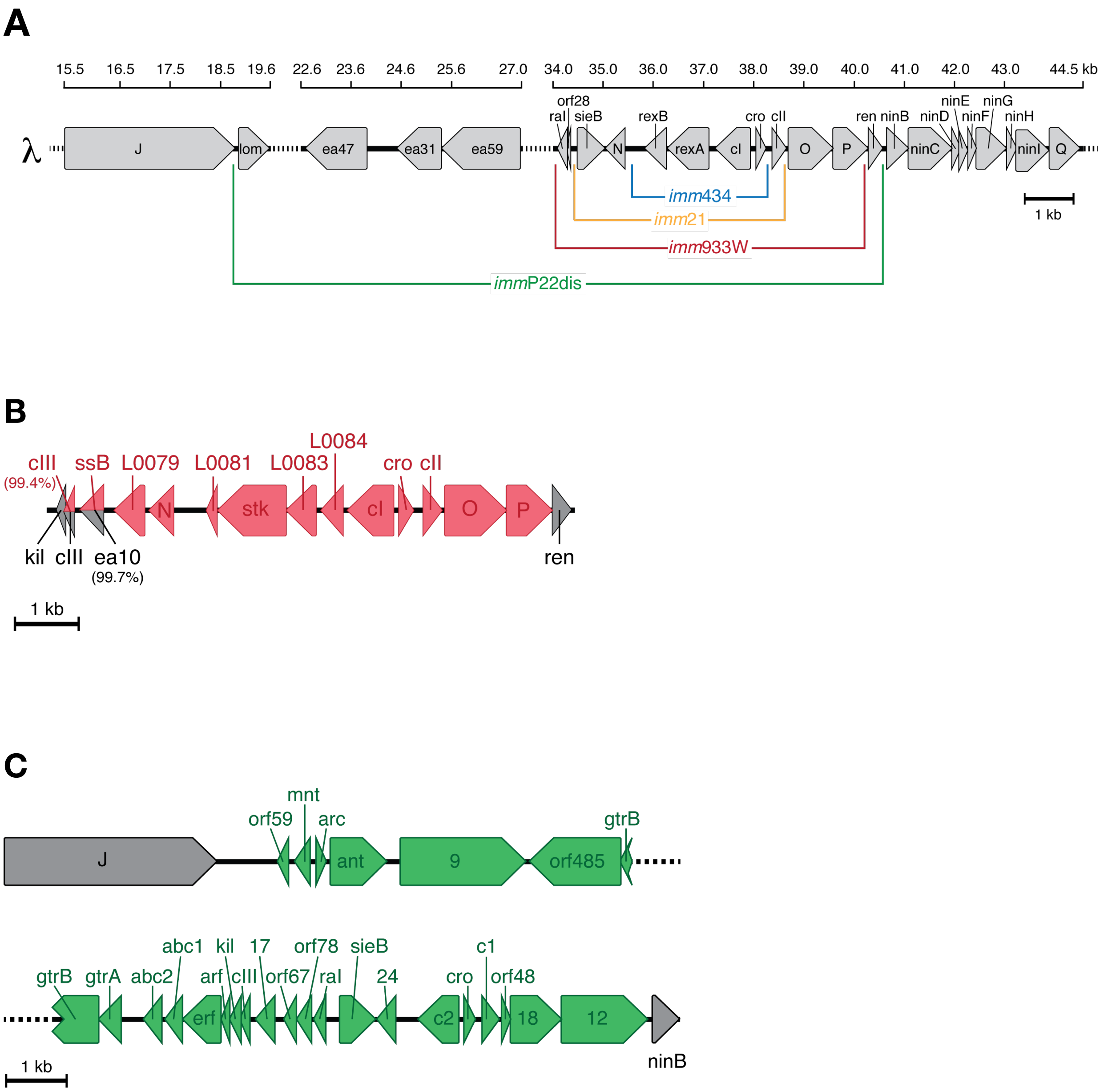

**Figure S3.** Genetic maps of λBH1 and λBH2 phage.

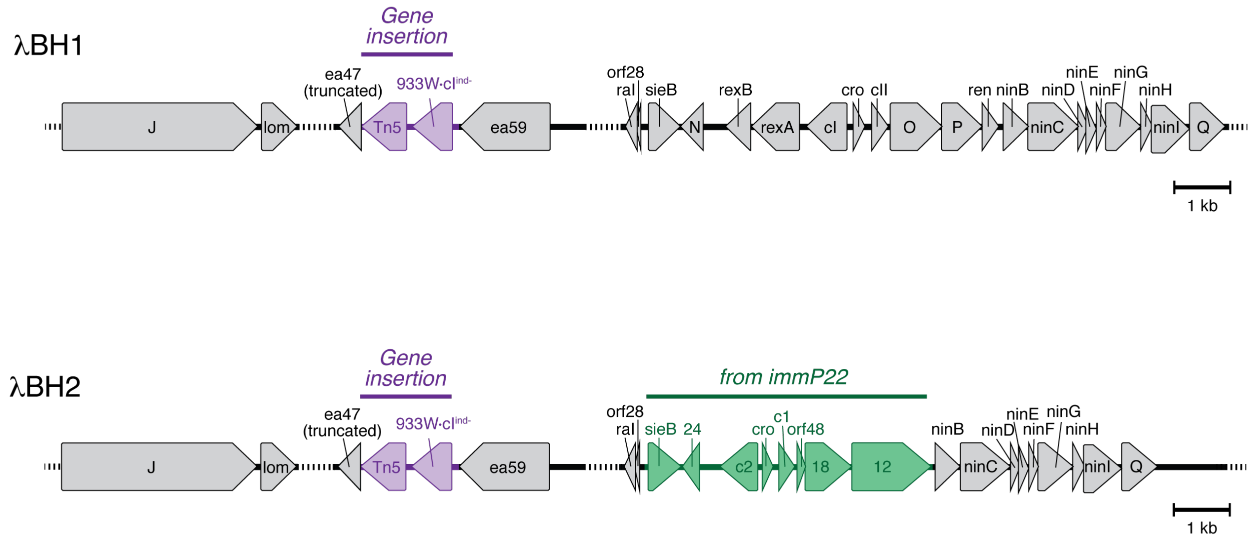

**Figure S4.** Spot assays of 3 μL of ~$10^7$ pfu/mL of λ, λ*imm*933W, λ*imm*434, and λ*imm*P22dis phages against non-lysogenic *E. coli*, its λ, λBH1, or λBH2 lysogen

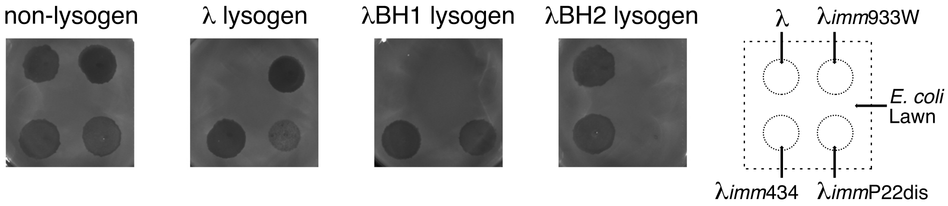

**Table S1**. Bacteria used in this study

| Bacteria | Description | Source |
|---|---|---|
| *E. coli* | Strain MG1655 with spontaneous streptomycin resistance | ATCC |
| *E. coli* C600 | Strain C600 | ATCC |
| *E. coli* $^{933W}$ | *E. coli* lysogenized by 933W phage derived from E. coli O157:H7 | This paper |
| *E. coli* O157:H7 | Strain edl933 | ATCC |
| λ*imm*933W lysogen | *E. coli* lysogenized by λ*imm*933W | This paper |
| λ lysogen | *E. coli* lysogenized by λ | This paper |

**Table S2**. Phages used in this study

| Phage | Description | Source |
|---|---|---|
| λ | Wild-type | ATCC |
| λ*imm*933W | | Gerald Koudelka |
| λ*imm*21 | | Donald Court |
| λ*imm*434 | | Donald Court |
| λ*imm*P22dis | | Donald Court |
| λBH1 | λ; ea47-*ea31::Tn5-933W·cI* $^{ind-}$ | This paper |
| λBH2 | λ; *ea47-ea31::Tn5-933W·cI* $^{ind-}$; *λ·sieB-λ·ren::P22·sieB- P22·12* | This paper |

**Table S3**. DNA sequences in this study

| Region | Sequence (5' to 3') |
|---|---|
| Tn5 and 933WcI $^{ind-}$ genes for insertion into λ. Flanking homology regions for ea59 and ea47 are shown in gray text. The LacUV5 promoter is underlined with mutated nucleotides (uppercase) for constitutive expression. The non-inducible repressor, 933W·cI $^{ind-}$ is shown in unformatted black text. A kanamycin resistance cassette (Tn5) is shown in blue text. | agaggttcatctgcatccacctttgctctctgctttttacgaacattaagcgacttactcgatgcacgca<br>atggtgtagcaataattgcaactcattccccagtagtactgcaagaggttccaaaatcctgcatgtggaaa<br>gtcctacggtcaagagaagcaataaatattatccgtccggatattgagacattcggtgagaacttaggtgt<br>tttaactcgtgaggtgtttttacttgaagtgacaaattctggataccaccacttattatcgcagtccgttg<br>attcagagctttcttatgaaaccattctaaaaaattataatggtcagataggattagaaggtcgaaccgtt<br>ttaaaagcgatgataatgaacagagatgaaggtaaagtacaatgagcgcaacgcaattaatgtgagttagc<br>tcactcattaggcaccccaggctttacactttatgcttccggctcgtatAAtgtgtggaattaTgagcgga<br>taaTaatttcacacaggaaacagctatggttcagaatgaaaaagtgcgcaaagaattcgcccagcggctag<br>cgcaagcctgtaaagaagctggtcttgatgaacatggtaggggaatggctatagcccgtgccctttctctt<br>tcgtccaaaggcgttagcaaatggtttaatgctgagtctttaccgcgtcaggaaaaaatgaatgcgcttgc<br>gaaatttctaaacgttgatgttgtttggcttcagcacggcacttcgttaaatggagcgaatgatgaagata<br>ctctttcatttgttggcaaattaaaaaaagggttagtgcgcgtggttggtgaggcaattcttggtgttgat<br>ggtgccatcgagatgaccgaagagcgcgatgggtggctcaaaatttatagcgatgatccagatgcctttgg<br>tcttcgtgtgaaaggagacagcatgtggcccagaataaaatcaggagaatatgtactcattgagcctaaca<br>ccaaagtattcccgggtgatgaggtgtttgtcagaaccgttgaaggacacaacatgattaacgttcttggc<br>tatgacagagatggagaataccaatttacaagcattaaccaggatcacaggcctataacgttgccttatca<br>tcaagtagcaaaggtggagtatgtagctggtattctgaagcaatctcgccatctggatgacatcgaggcaa<br>gggagtggctgaaaagttcgtgagccctgcaaagtaaactggatggctttcttgccgccaaggatctgatg<br>gcgcaggggatcaagatctgatcaagagacaggatgaggatcgtttcgcatgattgaacaagatggattgc<br>acgcaggttctccggccgcttgggtggagaggctattcggctatgactgggcacaacagacaatcggctgc<br>tctgatgccgccgtgttccggctgtcagcgcaggggcgcccggttctttttgtcaagaccgacctgtccgg<br>tgccctgaatgaactgcaggacgaggcagcgcggctatcgtggctggccacgacgggcgttccttgcgcag<br>ctgtgctcgacgttgtcactgaagcgggaagggactggctgctattgggcgaagtgccggggcaggatctc<br>ctgtcatctcaccttgctcctgccgagaaagtatccatcatggctgatgcaatgcggcggctgcatacgct<br>tgatccggctacctgcccattcgaccaccaagcgaaacatcgcatcgagcgagcacgtactcggatggaag<br>ccggtcttgtcgatcaggatgatctggacgaagagcatcaggggctcgcgccagccgaactgttcgccagg<br>ctcaaggcgcgcatgcccgacggcgaggatctcgtcgtgacccatggcgatgcctgcttgccgaatatcat<br>ggtggaaaatggccgcttttctggattcatcgactgtggccggctgggtgtggcggaccgctatcaggaca<br>tagcgttggctacccgtgatattgctgaagagcttggcggcgaatgggctgaccgcttcctcgtgctttac<br>ggtatcgccgctcccgattcgcagcgcatcgccttctatcgccttcttgacgagttcttctaaaattttt<br>atgataaacaattccatccaattgatttaatcaatacaacatttgaagatcaagcagataaatatatttt<br>tggcgttatgcagctgacagagccaaaataacaaatgcctatggcttcatttggatatcagagctatggct<br>cagaaaagcaagcatctactccaataaaccaatacatacaatgccaattatagatgaaagacttcaggtaa |

Region of 933W phge in λ*imm*933W phage (blue text) flanked by the *cIII* and *ren* of λ (black text).

```
ttggaattgattcaaataataatcaaaaatgtatttcatggaaaatagttagagaaaacgaagaaaaaaa
ccgactttagaaatatcaacagcagactcaaaacatgacgaaaaaccatatttcatgcgttcagtcttaaa
agcaattggcggtgatgtaaacactatgaacaattga
ttagtctggatagccataagtgtttgatccattctttgggactcctggctgattaagtatgtcgataaggc
gtttccatccgtcacgtaatttacgggtgattcgttcaagtaaagattcggaagggcagccagcaacaggc
caccctgcaatggcatattgcatggtgtgctccttatttatacataacgaaaaacgcctcgagtgaagcgt
tattggtatgcggtaaaaccgcactcaggcggccttgatagtcatatcatctgaatcaaatattcctgatg
tatcgatatcggtaattcttattccttcgctaccatccattgaaggccatccttcctgaccatttccatca
ttccagtcgaactcacacacaacaccatatgcatttaagtcgcttgaaattgctataagcagagcatgttg
cgccagcatgattaatacagcatttaatacagagccgtgtttattgagtcggtattcagagtctgaccaga
aattattaatctggtgaagtttttcctctgtcattacgtcatggtcgatttcaatttctattgatgctttc
cagtcgtaatcaatgatgtatttttgatgtttgacatctgttcatatcctcacagataaaaaatcgccct
cacattggagggcaaagaagatttccaataatcagaacaagtcggctcctgtttagttacgagcgacattg
ctccgtgtattcactcgttggaatgaatacacagtgcttattcgtactaataaaatacccaattttctgtt
tcttggttgtgtccaaagttatattcaatatctggtgttgatgtatcaatattcttcataccatcaacaag
agttgatacaacagccaaatcttgtttgattctcattaaatggtatttcttccggcgcaataaactttcaa
tggcaagtttcttcgttgggaatgcaaaagatctttctgcatttttttgctactttcttaattgcatatcta
tttctcctttgtttccattcctgtaaccactgatttggtgctggtttaaaattaacaatccaatgcgcagg
aaccaaccatgcataatgctctgtctgatgaaaagctatatattgaagtgcgaatattttttatcccatctt
cttcaactgtcgcctggaatctccagaaaacaggcattccatcatgttcagtttctgattcaggaaaaggt
acgctccatgatttttgtcatatctcacctcaaataagtggtttgctgcctaatttcatttttctggcgacca
acacaagtcacctcgccgtcagttgttttgatttccggtagcctgccgcgtaaatggctacgtttggaaga
catacaccagtttctggttgcttatgtccaaactcattcgcgtacacaatggccgctcgctccagattgcg
tctgtattctttctgttgccagatcacgtcctgtgccatgaacttaattggcttagcgtcttctatgcgct
caggcgtttcgtgagtacctttagcctgaatctgcgctctgcttagagtagggcggtgtaatacttctgaa
cttattgcttcttcgcgggccagtacgccgttagctaatgcctttgcctttaaacgctcacgacgacgaga
acgtgaattgcctttgaactgagttctgcgtgtcatatagacctcctgatgaactttggtggtgtggtagg
tgggagacccatttcgacctgtttcggcctacttcaattcggcaatagtcccgcaggcctcgccgctttac
gtgcgacatattcccgtccatgaacccttcaccacaccccaaagttcactttggttattgcgctttgtcag
cgccgtagattcatattcgaatcgttgtatattcaccgccctggtgagtaatgcgtcctgctgacgacgat
aataatgaaccaatagttcgacattatcaagaactattggtacgaattttggtgatttattaactctacga
agtatgattctgatatataaggaaatttatttttgaaaatgtggctgatgaaggttatgcggcagggatca
taactgcatggtttagcgagttacatcaataaatacaattggttatgtttttttaggtgggcgaacgtgagg
caaagaaaacccggcgctgaggccgggctagattttaaagtatttatcttttagagatgtagatgtaaaac
ttttcgcctttgaaaatttttttgtcatcagaagggcttatgaactcatctttttttgtagggaaccgctaat
gctgcatcacgtctgcgaggcagcttgcttacttcctcgcgctttttcatgatcagttatcctttaataac
ctatacagttttgtaggggtacatcctgaggatattgttaagttcgtagcacgccttttccgcccatcatc
gtataaacgaaaaccagtagtagacgaattttctgcgtcaaaaactatagacagtatagcgtccccagact
ttttttgccattcgcatgtgccgttagttggtttcgtcatctgtagacgccagtcaagaacgccatcactt
atagctgagagatcgtttagtacatctagtacggattgatatctttcatttggatctacatgaatgcattt
gttcactattgttattaatttttttatgtatatggggaggatactcttttaatggatagcagccattaatta
tcgactctctgagttgttcaatcgtgctaaatgcagatctttctctttcaaaattatcatgtccaacacac
attctatatatggttaatcctgcctgatatatgtcatatgtgaaattataatcatttgttgataaagaaaa
atattccggtggcacatgaaaatgatatccaaactcaggcgcagctctcgattcctcattgactaactgag
ataatccaaagtcagatagcatggcctcatttctgtttgatatcataatgttattaggttttatatcaaaa
tgcataagaccttttgagtgtatatgataaagtccacttaaaaattgaatggaataccgtattatctccct
gcttgtaagattattttttttcattaattggtttagcgaaccattatgataaaatggcatggctatataga
tattgctctcacattgagcagcatactgaacttgcacaatatttggatgtgcatgtttatagagaagcctt
gcttcattaaagtagtcgtcgtggttagtgttttctttttttttctatttctttaatcaccaagtcatgagc
taggtgtctgtcatgagccagatatacttttgaaaaacaaccctgttcttctagatcactaatccattcga
attctacatcagctcttttgtatggagttagcatccccttacctccgcagatagtgcagccaaaacagctt
catttgtttcagttgtaaaaccagaattatcgattccatttatattgcggtgtgacttcaatatttcttta
tactcgatctctgttaggttcaatgatgatttcatgccagattttctaatagtgtaatattttcttacatc
actgcttgaaaatgcttcttgaataacagcttctatataaaggcggtcaatgctaagattatcagagtttg
attcagtaacgcgtatggcagctaattcaacattatataaattaagaatgtcgaggatgttatttctcaca
tactttaatttttctggtgtgtctaaggtcgaaggtattttaataacatcaacacatttgagtgcagactc
attagtgcaatatacaacaaaagatgtaactttgggcgccgctctaacacctagtattctcatttttttata
tcctatttagaatcaggccgcatctctgcgaccatccatcatccaaacgtctcttcactcatccgaagaa
gcagcaatccgggttagcacgcacaagctcaagcgcatcagtcagcgaaagttcagtactgtactgatgcc
atttcatatccttccgcatccaatagattttccatctatccagagaacgtatgtacttgattcttgctgat
ggcaggatgtttgtttcacctggattgccctgccacacggggcgctgttcgccgatatctatcgtttggtc
attgatgctataaacaatatccagttcattgcggatatgttcaggcggccttatgctttcaatgaattggt
gaacttcttttttttactgcttgatattcaaggtcattgaacgccatctatcctccttacccaaacgtctct
tcaggccactggttaccagctatgtgacgatgaagtcacgaacttttcagccactcccttgcctcgatgtc
atccagatggcgagattgcttcagaataccagctacatactccacctttgctacttgatgataaggcaacg
ttataggcctgtgatcctggttaatgcttgtaaattggtattctccatctctgtcatagccaagaacctta
atcatgttgtgtccttcaacggttctgacaaacacctcatcacccgggaatactttggtgttaggctcaat
gagtacatattctcctgattttattctgggccacatgctgtctcctttcacacgaagaccaaaggcatctg
gatcatcgctataaatttttgagccacccatcgcgctcttcggtcatctcgatggcaccatcaacaccaaga
attgcctcaccaaccacgcgcactaaccctttttttaatttgccaacaaatgaaagagtatcttcatcatt
cgctccatttaacgaagtgccgtgctgaagccaaacaacatcaacgtttagaaatttcgcaagcgcattca
tttttcctgacgcggtaaagactcagcattaaaccatttgctaacgcctttggacgaaagagaaagggca
cgggctatagccattcccctaccatgttcatcaagaccagcttctttacaggcttgcgctagccgctgggc
```

Region of P22 phage in λ*imm*P22dis phage (blue text) flanked by the *J* and *ninB* genes of λ (black text).

gaattctttgcgcactttttcattctgaaccatgagtacgatactaaagcacttgcaaaaactttcagttc
aaccataatacgtactgaaagtacgaaaaaggatattcctatgcaaaatcttgatgagccgattaaaggtg
tcggcatccctgaagttgcgaaggcttgtggagttagcgaaagggctgtctataagtggctcaaaaacggc
ttcctccctaagactgagtttttgggaaaactaaatacgcatcaaaaatcgaagagatttctggtggcaa
atatcaagcaagcgaaatgcttgaaataagcaaaaagaaccttctggctgcataagtaacaccgctatttt
cacaatggacattcgtcctacgtcgctgacaaagcgagtcccaatatatctgaccaactaaggccatatgc
gtttccacgcatacctttcaactagctattcactattggaaatcttaagaaatggaacaaacaagttacag
caaactatcacagcgagaaattgatcgcgctgaaactgatttactcatcaacctgtcaacgcttacccagc
gcggtctggcaaagatgattggctgtcatgaatcgaagataagcagaacagactggaggtttatagcttcg
gtcttgtgtgcttttggcatggcatcagacatcagtccgattagcagagcttttaagtatgcgcttgatga
aatcacaaagaaaaaatccccggtggccgccggggactctaagcaaattgatatgcaattctgagggaatt
actggatcaatccacaggagtcattatgacaaatacagcaaaaatactcaacttcggcagaggtaactttg
ccgaacaggagcgtaatgtggcagatctcgatgatggttacgccagactatcaaatatgctgattgaggct
tattcaggcgcagatctgaccaagcgacagtttaaagtgctgcttgccattctgcgtaaaacctatgggtg
gaataaaccaatggacagaatcaccgattctcaacttagcgagattacaaagttacctgtcaaacggtgca
atgaagccaagttagaactcgtcagaatgaatattatcaagcagcaaggcggcatgtttggaccaaataaa
aacatctcagaatggtgcatccctcaaaacgagggaggttcccctaaaatgagggacatccctcaaaacga
gggaaaatcccctaaaacgagggataaaacatccctcaaattaggggattgctatccctcaaaacaggggg
acacaaaagacactattacaaaagaaaaaagaaaagattattcgtccgagaattctggcgaatcctctgac
cagccagaaaacgatctttctgtggttaaaccggatgctgcaattcagagcggcagcaagtggggaacagc
agaagacctgaccgccgcagagtggatgtttgacatggtgaagaccatcgcaccatcagccagaaaaccga
attttgcagggtgggctaacgatatccgcctgatgcgtgaacgtgacggacgtaaccaccgcgacatgtgc
gtgctgttccgctgggcatgccaggacaacttctggtccggtaacgtgctaagtccggccaaactccgcga
caagtggacccaactcgaaatcaaccgtaacaagcaacaggctggcgtgacagctggaaaaccaaaactcg
acctgacaaacactgactggatttacggggtggatttatgaaaaacatcgccgcacagatggttaactttg
accgtgagcagatgcgtcggatcaccaacaacatgccggaacagtacgacgaaaagccgcaggtacaacag
gtagcgcagatcatcaacggtgtgttcagccagttactggcaactttcccggcgagtctggctaaccggga
ccagaacgaactgaatgaaatccgccgccagtgggttctggctttccgggaaaacggatcacctcgatgg
aacaggttaacgcaggaatgcgcgtagcccgtcggcagaatcgaccatttcttccatcacccgggcagttt
gttgcatggtgccgggaagaagcatccgttatcgccggactgccaaacgtcagcgagctggttgatatggt
ttacgagtattgccggaagcgaggcctgtatccggatgcagagtcttatccgtggaaatcgaacgcgcact
actggctggttaccaacctgtaccagaacatgcgggccaatgcgctgactgacgcggaattacgacgcaag
gctgccgatgaactgacctgtatgacagcgcgaattaaccgtggtgagacgatacctgaaccagtaaaaca
acttcctgtcatgggcggtagacctctaaatcgtgcacaggctctggcgaagatcgcagaaatcaaagcta
agttcggactgaaaggagcaagtgtatgacgggcaaagaggcaattattcattacctggggacgcataata
gcttctgtgcgccggacgttgccgcgctaacaggcgcaacagtaaccagcataaatcaggccgcggctaaa
atggcacgggcaggtcttctggttatcgaaggtaaggtctggcgaacggtgtattaccggtttgctaccag
ggaagaacgggaaggaaagatgagcacgaacctggtttttaaggagtgtcgccagagtgccgcgatgaaac
gggtattggcggtatatggagttaaaagatga
atgggtaaaggaagcagtaaggggcataccccgcgcgaagcgaaggacaacctgaagtccacgcagttgct
gagtgtgatcgatgccatcagcgaagggccgattgaaggtccggtggatggcttaaaaagcgtgctgctga
acagtacgccggtgctggacactgaggggaataccaacatatccggtgtcacggtggtgttccgggctggt
gagcaggagcagactccgccggagggatttgaatcctccggctccgagacggtgctgggtacggaagtgaa
atatgacacgccgatcacccgcaccattacgtctgcaaacatcgaccgtctgcgctttaccttcggtgtac
aggcactggtggaaaccacctcaaagggtgacaggaatccgtcggaagtccgcctgctggttcagatacaa
cgtaacggtggctgggtgacggaaaaagacatcaccattaagggcaaaaccacctcgcagtatctggcctc
ggtggtgatgggtaacctgccgccgcgcccgtttaatatccggatgcgcaggatgacgccggacagcacca
cagaccagctgcagaacaaaacgctctggtcgtcatacactgaaatcatcgatgtgaaacagtgctacccg
aacacggcactggtcggcgtgcaggtggactcggagcagttcggcagccagcaggtgagccgtaattatca
tctgcgcgggcgtattctgcaggtgccgtcgaactataacccgcagacgcggcaatacagcggtatctggg
acggaacgtttaaaccggcatacagcaacaacatggcctggtgtctgtgggatatgctgacccatccgcgc
tacggcatggggaaacgtcttggtgcggcggatgtggataaatgggcgctgtatgtcatcggccagtactg
cgaccagtcagtgccggacggctttggcggcacggagccgcgcatcacctgtaatgcgtacctgaccacac
agcgtaaggcgtgggatgtgctcagcgatttctgctcggcgatgcgctgtatgccggtatggaacgggcag
acgctgacgttcgtgcaggaccgaccgtcggataagacgtggacctataaccgcagtaatgtggtgatgcc
ggatgatggcgcgccgttccgctacagcttcagcgccctgaaggaccgccataatgccgttgaggtgaact
ggattgacccgaacaacggctgggagacggcgacagagcttgttgaagatacgcaggccattgcccgttac
ggtcgtaatgttacgaagatggatgcctttggctgtaccagccgggggcaggcacaccgcgccgggctgtg
gctgattaaaacagaactgctggaaacgcagaccgtggatttcagcgtcggcgcagaaggcttcgccatg
taccgggcgatgttattgaaatctgcgatgatgactatgccggtatcagcaccggtggtcgtgtgctggcg
gtgaacagccagacccggacgctgacgctcgaccgtgaaatcacgctgccatcctccggtaccgcgctgat
aagcctggttgacggaagtggcaatccggtcagcgtggaggttcagtccgtcaccgacggcgtgaaggtaa
aagtgagccgtgttcctgacggtgttgctgaatacagcgtatgggagctgaagctgccgacgctgcgccag
cgactgttccgctgcgtgagtatccgtgagaacgacgacggcacgtatgccatcaccgccgtgcagcatgt
gccggaaaaagaggccatcgtggataacggggcgcactttgacggcgaacagagtggcacggtgaatggtg
tcacgccgccagcggtgcagcacctgaccgcagaagtcactgcagacagcggggaatatcaggtgctggcg
cgatgggacacaccgaaggtggtgaagggcgtgagtttcctgctccgtctgaccgtaacagcggacgacgg
cagtgagcggctggtcagcacggcccggacgacggaaaccacataccgcttcacgcaactggcgctgggga
actacaggctgacagtccgggcggtaaatgcgtgggggcagcagggcgatccggcgtcggtatcgttccgg
attgccgcaccggcagcaccgtcgaggattgagctgacgccgggctattttcagataaccgccacgccgca
tcttgccgtttatgacccgacggtacagtttgagttctggttctcggaaaagcagattgcggatatcagac
aggttgaaaccagcacgcgttatcttggtacggcgctgtactggatagccgccagtatcaatatcaaaccg
ggccatgattattactttatatccgcagtgtgaacaccgttggcaaatcggcattcgtggaggccgtcgg

tcgggcgagcgatgatgcggaaggttacctggatttttttcaaaggcaagataaccgaatcccatctcggca
aggagctgctggaaaaagtcgagctgacggaggataacgccagcagactggaggagttttcgaaagagtgg
aaggatgccagtgataagtggaatgccatgtgggctgtcaaaattgagcagaccaaagacggcaaacatta
tgtcgcgggtattggcctcagcatggaggacacggaggaaggcaaactgagccagtttctggttgccgcca
atcgtatcgcatttattgacccggcaaacgggaatgaaacgccgatgtttgtggcgcagggcaaccagata
ttcatgaacgacgtgttcctgaagcgcctgacggcccccaccattaccagcggcggcaatcctccggcctt
ttccctgacaccggacggaaagctgaccgctaaaaatgcggatatcagtggcagtgtgaatgcgaactccg
ggacgctcagtaatgtgacgatagctgaaaactgtacgataaacggtacgctgagggcggaaaaaatcgtc
ggggacattgtaaaggcggcgagcgcggcttttccgcgccagcgtgaaagcagtgtggactggccgtcagg
tacccgtactgtcaccgtgaccgatgaccatccttttgatcgccagatagtggtgcttccgctgacgtttc
gcggaagtaagcgtactgtcagcggcaggacaacgtattcgatgtgttatctgaaagtactgatgaacggt
gcggtgatttatgatggcgcggcgaacgaggcggtacaggtgttctcccgtattgttgacatgccagcggg
tcggggaaacgtgatcctgacgttcacgcttacgtccacacggcattcggcagatattccgccgtatacgt
ttgccagcgatgtgcaggttatggtgattaagaaacaggcgctgggcatcagcgtggtctgagtgtgttac
agaggttcgtccgggaacgggcgttttattataaaacagtgagaggtgaacgatgcgtaatgtgtgtattg
ccgttgctgtctttgccgcacttgcggtgacagtcactccggcccgtgcggaaggtggacatggtacgttt
acggtgggctattttcaagtgaaaccgggtacattgccgtcgttgtcgggcggggataccggtgtgagtca
tctgaaagggattaacgtgaagtaccgttatgagctgacggacagtgtgggggtgatggcttccctggggt
tcgccgcgtcgaaaaagagcagcacagtgatgaccggggaggatacgtttcactatgagagcctgcgtgga
cgttatgtgagcgtgatggccggaccggttttacaaatcagtaagcaggtcagtgcgtacgccatggccgg
agtggctcacagtcggtggtccggcagtacaatggattaccgtaagacggaaatcactcccgggtatatga
aagagacgaccactgccagggacgaaagtgcaatgcggcatacctcagtggcgtggagtgcaggtatacca
tgaaaaaagttaacattggaaacgtaccaaagatgctcgttccgctctttgagagcggtacaattgtgttt
tgtagagactttccagaatggcaacgcctgcatcaaaaacttggtgtggacgtgcaggactcggatgccaa
cggagcgtctcatacaatgagcagcgagaatggtgttttgcatgtgataggcgtgttcaatggcaaactat
ctactattgcccatgagtgcgctcacatggcattcgatatctgctcaagggtcggtgttgatgttgaacca
ggaagagccaacgagacttactgctacttaatgagcaggcttgttgagttctgcgagcgacatatcaaaaa
gccggagtgacccggcttgattattacttttttgctgtctggagttcgcttatctaatacccagccattacc
tggctttgttgttggtggaagcctttcgttgtccttgacggtggcaaaattgtctttcttaccgcctcgcg
ggccaacttcttggtatattccgccgtttttttcctgtgttttcacctggtttttttcgccatgatatacctc
aacatacacccgttattgggcgattaaatattgatctcattttataagtagtcaatatggcccaggtaaat
gcaaaaattaacccgccgtcaggtggttttttttgtacaaatccttcagcgtatcaaacaccatcttcttaa
caagctctgactgctcatcagcgagtcgttctgcatcgttgcgatatccagtcacaggcgatggttttgat
agagcatcttggacgatttgtaacaactcggagttcattgatctcccattcgcctccgccctgaattttaa
tttctccctgacttccataggcatacggaagttaaagtgcggatcatctctagccatgccatcactccaag
ttagtgtattgacatgatagaagcactctactatattctcaataggtccacggtggacctgtattgtgagg
tgaatatgaaaggaatgagcaaaatgccgcagttcaatttgcggtggcctagagaagtattggatttggta
cgcaaggtagcggaagagaatggtcggtctgttaattctgagatttatcagcgagtaatggaaagctttaa
gaaggaagggcgcattggcgcgtaaagttgaagccccaactgcggtaacagtcagggcttcggttgtcagt
aaatccttggagaaaaaccaacatgaatagtatagcaattttagaagcagttaacacctcttacgtgccgt
ttaatggacagcatgttcttaccgctatggtggctggagttgcctatgtagctatgaagccagtcgtggat
aacattggtctctcatggtcatctcaggtgcaaaagcttctgaaaatgaaagataaattcaactatgtcga
tatcgacatggttgctggagatatgaagaaacgtctcatgggatgcatcccactgaagaaacttaacggct
ggctgttcagcattaaccctgagaaagttcgtgcagacatccgtgacaaactgattaagtaccaggaagaa
tgcttcaccgttctgtatgattactggacgaaaggtaaggctgaaaacccgcgtaagaaaacatctgtcga
tgagaggacgccgcttcgtgatgctgtaaatatgcttgtaagcaaaaagcatctgatgtacccagaagctt
atgcaatgatccatcagcgtttcaatgtggaaagtattgaagaactggaggcgtctcagataccgctggcc
gtagagtacatccacagggtagtgcttgaaggtgagttcattggcaaacaagagaagaaaaccaacgatct
ttctgcaaaagaagcaaacagccttgtatggttatgggattatgccaaccgctcacaggcgttattccgcg
aactgtatcctgcaatgagacagattcaatctaactattcaggaaagtgctacgactacggccatgaattc
tcgtacatcattggaatagcgagagacgttttaattaatcacacgcgagatgttgatattaatgaacctga
cgggccaacgaatctttccgcatggatgagacttaaggataaagagcttccaccttcattacatcgctact
gacagataaccaacgcaacgacccagcttcggctgggttttttttatgcccaaaattcaccgtagccacgct
taggtaatgagcttgaaggagagacctacaaaaaaattgtaggtcgaaaagcgaacaaaataacttccgaa
aaagttgttttatcacaaaaaattcaccgtagccatgctgcggcaattccttgcatctggagcaaattaaa
tgacagacatcactgcaaacgtagttgtttctaaccctcgtccaatcttcactgaatcccgttcgtttaaa
gctgttgctaatgggaaaatttacattggtcagattgataccgatccggttaatcctgccaatcagatacc
cgtatacattgaaaatgaggatggctctcacgtccagattactcagccgctaattatcaacgcagccggta
aaatcgtatacaacggccaactggtgaaaattgtcaccgttcagggtcatagcatggctatctatgatgcc
aatggttctcaggttgactatattgctaacgtattgaagtacgatccagatcaatattcaatagaagctga
taaaaaatttaagtattcagtaaaattatcagattatccaacattgcaggatgcagcatctgctgcggttg
atggccttcttatcgatcgagattataatttttatggtggagagacagttgattttggcggaaaggttctg
actatagaatgtaaagctaaatttataggagatggaaatcttatttttacgaaattaggcaaaggttcccg
cattgccggggtttttatggaaagcactacaacaccatgggttatcaagccttggacggatgacaatcagt
ggctaacggatgccgcagcggtcgttgccactttaaaacaatctaaaactgatgggtatcagccaaccgta
agcgattacgttaaattcccaggaatagaaacgttactcccacctaatgcaaaagggcaaaacataacgtc
tacgttagaaattagagaatgtatagggtcgaagttcatcgggctagcggtctaatggctggtttttttgt
ttagagggtgtcacttctgcaagatggtagacgccaataatccaagcggaggtaaagatggcattataacc
ttcgaaaaccttagcggcgattgggggaagggtaactatgtcattggcggacgaaccagctatgggtcagt
aagtagcgcccagtttttacgtaataatggtggctttgaacgtgatggtggagttattgggtttacttcat
atcgcgctggggagagtggcgttaaaacttggcaaggtactgtgggctcgacaacctctcgcaactataat
ctgcaattccgcgactcggtcgttatttaccccgtatgggacggattcgatttaggtgctgacactgacat
gaatccggagttggacaggccaggggactaccctataacccaatacccactgcatcagttacccctaaatc

acctgattgataatcttctggttcgcggggcgttaggtgtaggttttggtatggatggtaagggcatgtat
gtgtctaatattaccgtagaagattgcgctgggtctggcgcgtacctactcacccacgaatcagtatttac
caatatagccataattgacaccaatactaaggatttccaggcgaatcagatttatatatctggggcttgcc
gtgtgaacggtttacgtttaattgggatccgctcaaccgatgggcagagtctaaccatagacgcccctaac
tctaccgtaagcggtataaccgggatggtagacccctctagaattaatgttgctaatttggcagaagaagg
gttaggtaatatccgcgctaatagtttcggctatgatagcgcagcgattaaactgcggattcataagttat
caaagacattagatagcggagcattgtactcccacattaacgggggggccggttctggctcagcgtatact
caacttactgctatttcaggtagcacacctgacgctgtatcattaaaagttaaccacaaagattgcagggg
ggcagagataccatttgttcctgacatcgcgtcagatgattttataaaggattcctcatgtttttttgccat
attgggaaaataattctacttctttaaaggctttagtgaaaaaacccaatggagaattagttagattaacc
ttggcaacactttagatatgtaataaaaatgggtgtaaacacccattttattttatgttaaatattctat
agctaattaaacctaacaactatggtttcccctacaacaccaatatcgtatacgttattaccagattttt
ccacccattttcaagtttaacctctttgtcatatagtctgtaatttctggaaaacacatttctttgcatta
acacctctgaccacatccaatcattgttaataatgcgtggtattaactctctcattaaaggatgctttatt
actatgttttcatttattggtgcatacggttctgtgccaatgaattttatattttttttgtctcttccaaa
tccaagataatctatgtcttgagatattctatttacaatgctttcctcaagctgaaactgtgcatttatgg
cattgtaagcaccataagaaaatattgttgatattaaaagaataaaagaaaaatatattcttgatattaac
tgtttatcttcaaaagcatagaatacgcataggcaacaaaaaaacataaagccacccataccaatcaatac
cctcggtgcgtatattggtgattttagaaaaatcattggtccaatgatgaagaacattgatgctaataaaa
ttaaaactactagcaataactttgttttcttattttcatctcttttgattgcttttaaaactatgactatc
aaagaaatgattagcgcaaagaatagcgagtagtagattaagtaattatcgccattcaagatcgtgctaaa
cattctataaaatgataagacgttagaaattatcccttcaaataaacttgagtttatctctataatcttac
tatgttcgatattgtaagaacctgttacaagtctttttgcaataaagtaagaataggcaaaatatcctact
attaaaccagcgacagaagatgctgtatttttttgtgatatttgaaattgagtttttcttaaccacatctga
aattataaaggccaacaagaatattgcgtaagtattcagcgcagcctgataaagactaaggaatgcaatgg
ttaaaatggatgatattatgatatttataggcttgtattgataagcgacatacgatgagataatagatatt
gccacactcatgcacattgttaatgaatcatatctatatgatagattttcaataaagaatgggtttgccaa
aatcatcataaaacaaagagatgctgtgatgtagtcatctccaaacagcttttccctgatgcaggatagtg
ccaatgctaaaataactatccctagcattaaaggtagcggagaagcatctataattggggttccaaaatta
atgatatagaaaataaagtcggaaagtgggcgaccattgcctgaccaacccaacccgccatataaagacct
acccaagtcatcaacgaaaaatgattgatgtgtcaataaaggaaatgtatatataatcgccaatccaagaa
agattgatataaatatcctgtcattactattaaatttcacttttaaaacccttacgctttaatatgtattt
aggccgctgtttggtttctatgtaaattctaccaatatattctccaagaatacctattcctatcaattgaa
cgccacccaggaaaagaacagaaacaagaagagacgggtagccaggaacattatttccaaatattaattta
tcaataatcatccatgcaccgtaaaggaatgacatacctgcaataaacaatccaatgtaagtccatatgcg
gagcggaaatgttgagaaagaagttattccctccagcgccaggttccataatttccagccgttgaatttcg
aatcaccggccacgcgttcggcacgggcatatttaacaacatccgtttttccgccaacccaactgagcaca
cccttcataaacaagttgcgttctggcatttgtttgatgttctcgacaaccgcacggctcattaaccgaaa
gtcgccaacattttcttcgatttttggattgctgattttattgtgcagcttataaaaccactcagctgtct
tacgcttcatgcgcccgtcagttgagcggtctgagcgcttagccagcaccatatccgcgccagcctgccac
ttctcaatgagatgagggataacttctatcggatcctgtaaatcgacatcaataggaatgaccgcatcccc
ggttgcatggtcgagacccgcgaaaagagcaggttctttaccgaagtttcgcgtaaacgaaagcggaataa
cgagcggatcagatgcagctattttgttaattattgattcagtcgcatctttactaccatcattaataaaa
acgatctcaatttcatattcttttagctcattaaactcacgtaccgttttatagaaaatcggtatcgtgtc
ttcttcgttaaaaactggaacgacaagagagattttcatcttatatccctgaaaacaatgaatctggaata
gataaagccgcataccaggctaattgccgagaaagtgataagggtaatcaatggtggcaaggaacattggt
cagccatccagccaacaacagcgctcagtgttcccatgaatcccacacacatcatgtagcgaagcgtggtg
gtggtggcattaaaggtgaaacgcgcattggcatagaagctgaacgatacggcgataacaaaaccggaaaa
gttcgccagcgcctgatgcgtatgcatcccatacacacaaaaagcaaatacgccccaatgaataagcgtgt
taagaacaccgatcgatgtgtacttagcgaataacttcaacattatgaaaatcagcggattcggaaaggtc
tgaagtgtagcactacaaattgttttgatcgatacaagcgatcaataatgtataatttgatagtttttatc
tatataatgcatgttaattgatcgttgttaccgatcaatttttattgctgattgctaagtggtttgggaca
aaaatgggacatacaaatctttgcatcggtttgcaaggctttgcatgtctttcgaagatgggacgtgtgag
cgcaggtatgacgtggtatgttgttgacttaaaaggtagttcttataattcgtaataataagttcggctgg
tagaatgtcgggattgtatgcaagtcctctcatcgtaaactcctcagttattgctgatagctccgtaacgc
gaacggtaatcacgaagacgcgggtctatttcaatgaatttggtgtaagtggctttgcggaatggccggat
ggctgtctggtaaattcgctcgcgttcttctttctctgcaagccatatacagtggcgaaattcctttcct
ctttcgtttcctgcggtagtgacattatcaggtcgtagtttttctgaatttatccagcacctccgatacg
gaattgccggaacagcggcgcgggtcatccgcaccatacaaaggcgctggcataatttactccagggtagg
ttatccgaataatgtggtacgtatagggttatttctttcgtaaacgtgatagcctgcttttaccgactct
tcacttcgcccgagaattttgctacatttctttgtgtatagcctgatgagataagcgtctgcattctttt
gtcttcgtcgtcgctccatcttggcttaatgaatgccgtttttaatgacagtttttttgctatgtaataaa
actgatttatgtttaggcccagatgttctgctgcacggcaagctaccatgcgaccgcaaactgactccatc
tccgctgtagttatgtttaatcttctcattaagccacctgtttaagctcatttattctgatattcattacc
tgaacgcatttgtctgctcatcatcgtgaccagtcaataattgccagtcgtgctggtatctctcaattag
ctttttcttgtccgtttctgttgctgcatattcactgaatgccttaagaacctgttcaggagcaggggagg
aaggcgatgatttagtttgcttagcaggtgctgcattctgctgctgtttgtgctcctcagtatcagcgtct
ttggcgtcgtcgataccaaacaaaccgttaaggcaatatttgcgagcgtaagagcttgtagcgcccgttac
ctgagctgcatccattcccttcttgttttcttcttctcgcgctatagcgcttgctgaatggctattttcac
catctgtaatggtcgctgtggccttgacgtaataacggtcgccaatcagcacgatttcatcactgatagac
aggaacagacctttcagtagtggcttaacaccctcaagaatgtcctcacaactgcggtatttgtatttacc
aaacgagttgtactgattctttggcgcattcagatgctcctgaatttcagcaagtcttgcgtaaaactctt
tgctcatgagtaataccccgcaaattcatcccaaccaataatcggattttgccgttctgcggctaagttga

tttgttgctcaacctcttcctcaatttcaggagagataagcgcgataaattcttcgtcactaaattcatgc
tgcatgatttcgattccagtcttcgtcctgacaatcttcccagcccattgcgattgatgatgcccatgcgt
atgcggcgctgttaccttccttcgtatccgggaaggatgcttcgtagagcttgttaaactcacgattgcct
tgctgcacaaggatggttccgttaacaggcacaatagtcatggctcggcactccaggctgattaaggatgt
ctgccagccgtttccagccagcgcgtaatttgcgggtgatgcgatctaaaagtgattcgtgtaactgggaa
acgcccatgcgagcgcttcccgcgattgcgataatcatgggagttccttatgttgtgtgtgattgcatgag
gctgagcacttgaataaatactcactcagatgcggatatgaaaaagccgcactcaggcggctgtcgtttct
tctttcaggctttcgagatattcacgcgggtcgtcgtaacactggcactcgctataccaatcaatccagcg
atcatccagttccatatcttccaaatcctggtcggtaaggctctcgtcaaacatctgtaaaccgttggcgt
tgcagtaatcaggcttgatgttgttgtcatactgaaaggcgtcataatcagccagtgcatccatcattcgt
acaccttcttcaacactacccacttctacaatgaatggcttcatagggacttgcgggatatgccagacacg
taatttcatatttcctccaggcaaaaagaatgccgcccatatagagcggcaagactatcaagggatgattc
tccaataaccagaacgagtcttcgtcctcattcggttacgagcgatattgctcacatagcagactcgtaaa
tctgctataggtgcttattcgcttggtggttcaggtaatggcatccagtgagtcactcgctcatgcttgat
taaacgcgtgtcctgctttgagttctttgcctcaattactggcttcctcatggtgtaaattcccgagaaat
ttgaattgcaaacacaatatccaagcacagtaattccggattccggcatccgctcactacacttaatccac
tccatcactccttccccagagccttgccgatggctgcgcgagccgtcgcgtacacagcgtcccactcgctc
acgtcatgttccatatcgataatgcttaacagagcctcgagaagctcaggagctgcacttgccaattttat
ttcctgcttattggcgttctcggctgaaaacagaatgtgaccttcagtaacatcgacaatgtaatcttcat
caacatcccaatgcggtatgtctttcatattcacctctgtggcttgctgccaaaagaagacagactatata
gcctttagtttttccagctctctggcaatcatttccgtggttctgattgcccatttatcgacaatctttcc
atcttctctcaccagagccatttcctcaggctttaccatacattcagcatcaagcttgcagcctttgcatt
tcacaaaacgactacaccattgatttgtatcaatagtcgtagtcatatgggtagtcctggtattgttccat
cacatcctgaggatgctcttcgaactcttcaaattcttcttccatatctcatctcaaatagtggattgcgg
tagtaaagattgtgcctgtcttttaaccacgtcaggctcggtggttctcgtgtacccctacagcgagaaat
cggataaactctattcaccccctacagagagtaaaaagagaatcgccgatgaacaactcatggtggcaggag
ttaatgcgtttttcctgcaaggaatgacacttaaacagttgattcatatgctaatcatcctgatcgtatt
gattattgttatgccggtaagcgtaaaagaatggataaacctgcataatccagaaatacttcctcattact
ggatgtattacatcctgttgttctgcgttagctatgtgcttaacggtgttgttaattccgtttatcacgct
gttactgaaagaattgaggcatcaactgctcagcggcgtaaggacagagaagaaaaagtcgttcggattt
gtttgattcgttaactcttggagaaagagcgtatttggcattcgctgtagccgctaataaccagctaaaga
cagaaaagggaagccctgaagcaatttcattgctcaaaaaagggattatcactcgattgccttctgctatt
ggatatcctgatattgaccgttttattatcccggaaaagtattttaatgagtgctacatgagatttgccgg
gaagtcagacattcttatgaatgaacttattgtacaggacgaacagctcaaaaaataacgacttaaccgac
aaataccttacctcgctgttatttgtttgctcttacgatgaccagccgcgtaaagtgctacgcctggaaga
agtacagatcctccttcaacttccttctgacgcgttccggcaagcgaaatggctttggtgacacggtcaat
tcttttggctttaacttcctgagaagcatcaggagcatcgcagccaaaaattgaatcgatgatattgcaga
tggtgtcgcgctctatggctagctttctgcgccgctcatgacggcgagttttagcattgcctgcaaacgtt
gacttcccgtaggtgataaccgtcatgatttaatcctcatgtgaaatggctttggtactggcgccggaacc
tgtctcaatttccggatttcaagtggcttctcagtccggcccgatcggtacagctagaggcctaagctcca
ccacacgccagtccaaaccaatctcgtttggtatttgttcgcgctttgtcagcgcatcatcgaagttaaag
agcgttgcctttccgtttggctaccagcgtcctgctgatggctaaaatttaagacttcttaattaaatggt
caagtgtatttttgaagaaaacttaaatattttatcgttacttaagtttttatttgatttttaaaggaaaa
tgtagtgtgagggggcgggtgccccttatggaagatttgcgagttttgcgtcaacaactacgccaatgattt
tgcagtttccgttgatttctatcatcggatattgtgggtttaatggttttaaaaactttcggcctgcatcc
ataactaatttttttgaatgtggcctcgttttcaccttctaattttgcaacaaccagcttgccgtttcttgg
ttcgacttcgggatcaaccagaattatcattccttctggaatgcttaaccctgccggtgctgtcatagagt
caccttggacatcaagccaaaatgaatcttctgaacaatctacagtggtgtcgtgccagttctctatcgcg
cgcttgtgataaggttctacagcttccatccattgccctgcgcttacccaactgataagagggtatgatcc
tcttggctcatgcctactatgataggcaacgtttgtctggcttaaatctcctttcagcaaatagtcagggg
agcactgaagagccttcgaaagtgccaacaggttctccccatttggctcagtctccgagcgctcccattgc
gatattgcaacattagacactcccaccatcttaccaagagcggcttgtctaatcttgagtttttttcttcg
agcgcgaatacgctcacccatcaattgtgtattcatagttaagtcatcttaaataaacttgactaaagatt
cctttagtagataatttaagtgttctttaatttcggagcgagtctatgtacaagaaagatgttatcgacca
cttcggaacccagcgtgcagtagctaaggctttaggcattagcgatgcagcggtctctcagtggaaggaag
ttatcccagagaaagacgcataccgattagagatcgttacagctggcgccctgaagtaccaagaaaacgct
tatcgccaagcggcgtaagcaaaacgctctttaccaatctgaaccgccgacaacgcggtaaacctatttca
aagcgcatcaacgaatgcgcacaactaactattaactacaggaatgttcacatatggaactcacaagcact
cgcaagaaagccaacgcaattaccagcagcatccttaaccggatagctattcgtggacagcgtaaagtcgc
tgatgcgttaggcattaacgaatctcaaatttcacgatggaaaggcgatttcattccgaagatggggatgt
tattggcggttctggagtggggtgtcgaggatgaggagttggcagaactggcaaagaaagttgcgcatctg
ctgacaaaagaaaagcctcaagactgcgggaacagttttgaggcctgatgtagaaagactggatcaatcca
caggagtaattatgccaaaacaactcagtcctgaccaggacaaattacacaaaaacatactacgtgatcgg
ttcttatccagcttcaaacagcctggtcgatttcgggctgagttggagaaagtgaagctaatactgaagag
gaaaggtcatgagtaatcttgcaacagttacaccgataaaacctcatctggaggttgtggagcatcgcgtg
gcagaactcgacgatggctacacccggactgcaaatacactgctggaagctgtcatgctttctgggcttac
tcaacatcagctactgattgttatggctgtgtggcgcaagacatacggttataacaaaaaaatagattgga
tcggaaatgaacagttcgctgaactcactggcatggcgccaaccaaatgttctaccgccaaaaacgagctt
atcagaatgggggttctcactcaggtggggcgtcaggttggtatgaataaaaatatttccgagtggaagac
gaaggttaacggattcggtaaaacatttaccagatcggtaaaactaaccttcaccaaatcggtaaaaacca
atttaccgaatcagtcaaacacaaaagacaatatacaaaagacaataaatacaaatacccccttacccct
aaagggggatgcgatgaaggttctaaacctgaaaagcgaaaacctaccaagattaactacagcgaatatct
tgctgcctacaacgagattgttggtgacagactcccacatgcagtggaggtcaattctgaacgacaacgca

Region of 21 phage in λ*imm*21 phage (blue text) flanked by the *orf28* and *O* genes of λ (black text).

```
agttgaaaaagctgattgattcactggcaaccaaaaacatcgacggattccgggcatacgtcaaagcgttc
atggcagcagccagaccattccatttcggtgataacgaccgtgactgggtagctaattttgattatctgct
acgcccgaaagtactgatagcaattcgtgagggaacactatgagacaggatatcgaggcgagcgttatcgg
tggcttgctgattggcggattaacaccaaccgccagtgacgttctggcaacactggagcctgaagcattct
caattccgctctaccggaaagcttttgaagttattcgaaagcaggccagaaacaggaacctgattgatgga
ctgatggtggccgaggagtgcggggatgaatacgcaacggcggtgatgatgactgcgcggtcatgtcccag
cgctgcaaacctgaaaggttatgccggaatggttgcagacagttatcaacggcgtcaggttttacagctac
tggatgagatgcgggagccaatcagtaacggcacgctggacgcatcaggcagagcgatggacgagcttgta
aagcgcctgtcatccatcaggaagccgcggaacgaggttaaacctgtgcgactgggtgaaatcatcaatga
ctacactgacacgcttgacaggcgtctgaggaacggagaagagtcggataccctgaagaccggaatcgaag
agcttgacgctatcaccggagggatgaacgcagaagaccttgtgattattgctgctcgtccaggtatgggt
aaaaccgaactggcgctgaagatagccgaaggcgtggcaagtcgtgttattcctggttctggcgtccggcg
cggtgtgttgatttctcgatggaaatgagcgccattcaggttgttgagagagggattgccggcgcaggaa
tgatgtcggtcagtgtgctgcgtaacccgtcacgtatggacgatgaaggatgggcgagagttgcaagcggg
atgaagttgctggcagagctggatgtgtgggtagttgacgcatcgcgtttgtctgtcgaagaaatcaggtc
catttccgaacgccacaagcaggagcatcctaatctgtcactgattatggctgactatctcgggctaattg
agaaaccaaaagcggaacgtaatgacctcgccatagcacatatctccggtagcctgaaagcgatggcgaaa
gacctgaaaactccagttatctccctaagccagctctcccgcgatgttgagaagcggccaaacaagcgccc
gacaaacgcagatttgcgggattcaggaagcattgaacaggacgcagactcaatcatcatgctctatcggg
aagcggtatatgacgagaacagtagcgccgcgccatttgctgaaatcatcgtgacgaaaaaccgttttggc
tcgcttggtacggtttaccagcggttctgcaacggacactttgttgcatgtgaccaggacgaagccagaca
gatttgcacggcatcaaatgcacctgctggacgcagaaagcgatatgcacaaggggctgacgtatgactat
ttacatcactgagttggtaacaggcctgctggtaatcgcaggccttttttatttggggggagagggaagtcat
gaaaaaactaacctttgaaattcgatctccagcacatcagcaaaacgctattcacgcagtacagcaaatcc
ttccagacccaaccaaaccaatcgtagtaaccattcaggaacgcaaccgcagcttagaccaaaacaggaag
ctatgggcctgcttaggtgacgtctctcgtcaggttgaatggcatggtcgctggctggatgcagaaagctg
gaagtgtgtgtttaccgcagcattaaagcagcaggatgttgttcctaaccttgccgggaatggctttgtgg
taataggccagtcaaccagcaggatgcgtgtaggcgaatttgcggagctattagagcttatacaggcattc
ggtacagagcgtggcgttaagtggtcagacgaagcgagactggctctggagtggaaagcgagatggggaga
cagggctgcatga
tcaatagtcgtagtcatacggatagtcctggtattgttccatcacatcctgaggatgctcttcgaactctt
caaattcttcttccatatatcacctcaaataagtggtttgctgcctaatttaattttctggcgaccaacac
aagtcacacccatttcactgcgtggcttgctgtagtaaatacggttctgtttacgctcgacttcttctgcc
ttcttgcagcgaaggcttccgagtgatgctgctttatctgctctgacgcaaccagagagctttagcgcaat
ttttcgcgccagtgcttcattactgcgtcgctcggcaataagttctgctctgcgagctttgtagcggcttt
ttgccgtacctttggattctttccagacaatggttaccatgatggtctcctttaagtggctttggcgcatg
acgcgtcgaggtgcttatcttctcgatcgctgtcttgtagctgcaattcgcgccatccccaaaaccactca
agttctggtctcaacggttaggttgagagtccgtcgatgttaaagagcctgccaatctgttccgtttggct
tccagcgtcctgctgatggcttaaatttaagacttcttaatttattggtcaagtgcatttttgaagaaaac
ttaattttatgggcgtgaatttagtttgtctttgatttttaacgggaaataaaaaaggggcgaaagcccct
taaggaaggtttgctagcttggcatcaacgacaacgccaatgattttacagttcccattgatttcaatcat
tgggtattgtggattgagtggtttcaggaattttctaccggcatcaataactaacttttgaatgtcgcct
cgttttctccttcaagtttggcgactaccagctttccattacgtggttcgacttctgggtcgacgagaata
atcatcccctcaggaatactcagtcctgccggggcagtcattgaatcgcctttaacgtcgagccaaaaaga
gtcttcagaacaatctaccgttgtgtcgtaccagttatctattgcacgcctatgatatggctctacagctt
ccatccaacatcctgcgcttacccaactaattagaggatacgaacctcttggatcatgcctgctgtgatag
gcaatgtttgaaagactatcctctcctttcaacaggtaatcaggggagcactgcaaagccttggctaaggc
caataggttttcgccattgggctcagtttcagatcgctcccattgggaaatagcaacattagacacgccaa
ccatcttgccaagggcagcctgcctaatcttgagttcttttctgcgagcgcgaatacgctcacccatcagt
tgtgtattcatagttaagacatcttaaataaacttgacttaagattcctttggtggataatttaagtgttc
tttaatttcggagcgagtctatgtacaaaaaagatgttattgaccacttcggaacccagcgtgctgttgct
aaagcactaggcattagcgatgcagcagtctctcagtggaaagaagttatcccagagaaagacgcctatcg
attggaaatcgttacagctggcgccctgaagtatcaagaaagtgcttaccgccaagcggcataagcaaatt
gctctttaacagttctggcctttcacctctaaccgggtgagcaaacatcagcggcaaatccattgggtgtg
ccgctataactcaatatcaatataggtaaattaacaaatggcacaagcaagctacagcaagccaacacagc
gagaaattgatcgcgctgaaactgatttactcatcaacctgtcaacgcttacccagcgcggtctggcaaag
atgattggctgtcatgaatcgaagataagcagaacggactggagatttattgcttcggtcttgtgtgcttt
cggaatggcatcagacatcagtccgattagcagggcttttaagtatgcgcttgatggactcacaaagaaaa
aacgcccggtgtgcaagaccgagcgttctgaacaaatccagatggagttctgaggtcattactggatctat
caacaggagtcattatgacaaatacagcaaaaatactcaacttcggcagaggtaactttgccggacaggag
cgtaatgtggcagatctcgatgatggttacgccagactatcaaatatgctgcttgaggcttattcgggcgc
agatctgaccaagcgacagtttaaagtgctgcttgccattctgcgtaaaacctatgggtggaataaaccaa
tggacagaatcaccgattctcaacttagcgagattacaaagttacctgtcaaacggtgcaatgaagccaag
ttagaactcgtcagaatgaatattatcaagcagcaaggcggcatgtttggaccaaataaaaacatctcaga
atggtgtatccctcaaaacgagggaaaatcccctaaaacgagggataaaacatccctcaaattggggggatt
gctatccctcaaaacagggggacacaaaagacactattacaaaagaaaaaagaaaagattattcgtccgag
aattctggcgaatcctctgaccagccagaaaacgatctttctgtggttaaaycggatgctgcaattcagag
cggcagcaagtggggaacagcagaagacctgaccgccgcagagtggatgtttgacatggtgaagaccatcg
cgccatcagccagaaaaccgaattttgctgggtgggctaacgatatccgcctgatgcgtgaacgtgacgga
cgtaaccaccgcgatatgtgtgtgcttttccgctgggcctgccaggacaacttctggtccggtaacgtgct
gagtccggccaaactccgcgacaagtggacccagctcgaaatcaaccgtaacaagcaacaggcaggcgtga
cagccagcaaaccaaaactcgacctgacaaacactgactggatttacggggtggatttatga
```

Region of 434 phage in λ*imm*434 phage (blue text) flanked by the *N* and *cII* genes of λ (black text).

atgatcgttatctgggttggacttctgcttttaagcccagataactggcctgaatatgttaatgagagaat
cggtattcctcatgtgtggcatgttttcgtctttgctcttgcattttcgctagcaattaatgtgcatcgat
tatcagctattgccagcgccagatataagcgatttaagctaagaaaacgcattaagatgcaaaacgataaa
gtgcgatcagtaattcaaaaccttacagaagagcaatctatggttttgtgcgcagcccttaatgaaggcag
gaagtatgtggttacatcaaaacaattcccatacattagtgagttgattgagcttggtgtgttgaacaaaa
ctttttcccgatggaatgggaagcatatattattccctattgaggatatttactggactgaattagttgcc
agctatgatccatataatattgagataaagccaaggccaatatctaagtaactagataagaggaatcgatt
ttcccttaattttctggcgtccactgcatgttatgccgcgttcgccaggcttgctgtaccatgtgcgctga
ttcttgcgctcaatacgttgcaggttgctttcaatctgtttgtggtattcagccagcactgtaaagtctat
cggatttagtgcgctttctactcgtgatttcggtttgcgattcagcgagagaatagggcggttaactggtt
ttgcgcttaccccaaccaacaggggatttgctgctttccattgagcctgtttctctgcgcgacgttcgcgg
cggcgtgtttgtgcatccatctggattctcctgtcagttagctttggtggtgtgtggcagttgtagtcctg
aacgaaaacaccccgcaatggcacattggcagctaatccggattcgcacttccggccaatgcttcgtttcg
tatcacacacaccaaagccttctgctttgaatgctgcccttcttcagggcttaattttaagagcctcacc
ttcaatggtggtcagtgcgtcctgctgatggcttaaaattacaaggaagattgtatgttgtaaacaataaa
tattgtaaaaaggggcgtgaaaaacaaactccattgttttaaacggaaaatagtttgtttttttgttatc
gagattgaggtggggattactgattgcaggttccgactacatcaccaacaaaggatttggttgatgtaagt
tgttgcatacctgggatgttcattactttggagtaaagagctttttttgtctgtagtgattgaccaagtttc
aacggttatgcctcctccagactggtattctcctaccatagtgttcgatgacaaagcagtgtatttcatct
ctggatagacgccagaaactgattcataaactgatgatttatcgccatttattgttacgttgaaaacggaa
tcttccgtgctgtcttttgtaaactcgtaacgatcgccattcattgccccgtacccgtgcaggtttgtgac
aatccagcattcagaattggcgctggtagttaagagtattgagagtagcgccgcaatcctgatcatacgaa
ttttaccctcgcttccacgacaacaccgataatcttgcagttcccgttgataggagtcataggccatgaag
gattcaggcctttcaggtacttctgaccgccatctatgaccagtttcttgaatgttgcttcgttcgcgtca
gtcagtttggctacaacaaggcttccattcactggctcgcgtccagtatctactaacaccatatgaccttc
agggatgctttgacctacaggtgaggtcatggaatcaccttcaaccttcagccagaatccattgcctaata
agttaacgtcactgtcataccattcatcaatgtccttgatatcgtagggttcacaagcttcacaccacgaa
ccagctctaaccatgctaatcaatggatatttccctttgggctcaacgtgcccaacaaatctaacattcga
atcagaggtgccattgagcagccagtcaacacttacgccaagagctgacgcaagttctggtaaaaagcgtg
gtcgcttagttttaccgttttcgagctgctctatagactgctgggtagtccccacctttttgagcaagttca
gcctggttaagtccaagctgaattcttttgctttttaccctggaagaaatactcataagccacctctgtta
tttaccccccaatcttcacaagaaaaactgtatttgacaaacaagatacattgtatgaaaatacaagaaagt
ttgttgatggaggcgatatgcaaactctttctgaacgcctcaagaagaggcgaattgcgttaaaaatgacg
caaaccgaactggcaaccaaagccggtgttaaacagcaatcaattcaactgattgaagctggagtaaccaa
gcgaccgcgcttcttgtttgagattgctatggcgcttaactgtgatccggtttggttacagtacggaacta
aacgcggtaaagccgcttaagacattcccgctcttacacattccagccctgaaaaagggcatcaaattaaa
ccacacctatggtgtatgcatttatttgcatacattcaatcaattgttatctaaggaaatacttacatatg
gttcgtgcaaacaaacgcaacgaggctctacgaatcgagagtgcgttgcttaacaaaatcgcaatgcttgg
aactgagaagacagcggaagctgtgggcgttgataagtcgcagatcagcaggtggaagagggactggattc
caaagttctcaatgctgcttgctgttcttgaatggggggtcgttgacgacgacatggctcgattggcgcga
caagttgctgcgattctcaccaataaaaaacgcccggcggcaaccgagcgttctgaacaaatccagatgga
gttctga

Region of P22 phage in λBH2 phage (blue text) flanked by the *orf28* and *ninB* genes of λ (black text).

tcaatagtcgtagtcatacggatagtcctggtattgttccatcacatcctgaggatgctcttcgaactctt
caaattcttcttccatatctcatctcaaatagtggattgcggtagtaaagattgtgcctgtcttttaacca
cgtcaggctcggtggttctcgtgtaccccctacagcgagaaatcggataaactctattcaccccctacagaga
gtaaaaagagaatcgccgatgaacaactcatggtggcaggagttaatgcgtttttttcctgcaaggaatgac
acttaaacagttgattcatatgctaatcatcctgatcgtattgattattgttatgccggtaagcgtaaaag
aatggataaacctgcataatccagaaatacttcctcattactggatgtattacatcctgttgttctgcgtt
agctatgtgcttaacggtgttgttaattccgtttatcacgctgttactgaaagaattgaggcatcaactgc
tcagcggcgtaaggacagagaagaaaaagtcgttcgggatttgtttgattcgttaactcttggagaaagag
cgtatttggcattcgctgtagccgctaataaccagctaaagacagaaaagggaagccctgaagcaatttca
ttgctcaaaaaagggattatcactcgattgccttctgctattggatatcctgatattgaccgttttattat
cccggaaaagtattttaatgagtgctacatgagatttgccgggaagtcagacattcttatgaatgaactta
ttgtacaggacgaacagctcaaaaaataacgacttaaccgacaaataccttacctcgctgttatttgtttg
ctcttacgatgaccagccgcgtaaagtgctacgcctggaagaagtacagatcctccttcaacttccttctg
acgcgttccggcaagcgaaatggctttggtgacacggtcaattcttttggctttaacttcctgagaagcat
caggagcatcgcagccaaaaattgaatcgatgatattgcagatggtgtcgcgctctatggctagctttctg
cgccgctcatgacggcgagttttagcattgcctgcaaacgttgacttcccgtaggtgataaccgtcatgat
ttaatcctcatgtgaaatggctttggtactggcgccggaacctgtctcaatttccggatttcaagtggctt
ctcagtccggcccgatcggtacagctagaggcctaagctccaccacacgccagtccaaaccaatctcgttt
ggtatttgttcgcgctttgtcagcgcatcatcgaagttaaagagcgttgcctttccgtttggctaccagcg
tcctgctgatggctaaaatttaagacttcttaattaaatggtcaagtgtatttttgaagaaacttaaata
ttttatcgttacttaagtttttatttgatttttaaaggaaaatgtagtgtgaggggcgggtgccccttatg
gaagatttgcgagttttgcgtcaacaactacgccaatgattttgcagtttccgttgatttctatcatcgga
tattgtgggtttaatggttttaaaaactttcggcctgcatccataactaatttttttgaatgtggcctcgtt
ttcaccttctaatttgcaacaaccagcttgccgtttcttggttcgacttcgggatcaaccagaattatca
ttccttctggaatgcttaaccctgccggtgctgtcatagagtcaccttggacatcaagccaaaatgaatct
tctgaacaatctacagtggtgtcgtgccagttctctatcgcgcgcttgtgataaggttctacagcttccat
ccattgccctgcgcttacccaactgataagagggtatgatcctcttggctcatgcctactatgataggcaa
cgtttgtctggcttaaatctcctttcagcaaatagtcaggggagcactgaagagccttcgaaagtgccaac
aggttctccccatttggctcagtctccgagcgctcccattgcgatattgcaacattagacactcccaccat
cttaccaagagcggcttgtctaatcttgagtttttttcttcgagcgcgaatacgctcacccatcaattgtg
tattcatagttaagtcatcttaaataaacttgactaaagattcctttagtagataatttaagtgttcttta

atttcggagcgagtctatgtacaagaaagatgttatcgaccacttcggaacccagcgtgcagtagctaagg
ctttaggcattagcgatgcagcggtctctcagtggaaggaagttatcccagagaaagacgcataccgatta
gagatcgttacagctggcgccctgaagtaccaagaaaacgcttatcgccaagcggcgtaagcaaaacgctc
tttaccaatctgaaccgccgacaacgcggtaaacctatttcaaagcgcatcaacgaatgcgcacaactaac
tattaactacaggaatgttcacatatggaactcacaagcactcgcaagaaagccaacgcaattaccagcag
catccttaaccggatagctattcgtggacagcgtaaagtcgctgatgcgttaggcattaacgaatctcaaa
tttcacgatggaaaggcgatttcattccgaagatggggatgttattggcggttctggagtggggtgtcgag
gatgaggagttggcagaactggcaaagaaagttgcgcatctgctgacaaaagaaaagcctcaagactgcgg
gaacagttttgaggcctgatgtagaaagactggatcaatccacaggagtaattatgccaaaacaactcagt
cctgaccaggacaaattacacaaaaacatactacgtgatcggttcttatccagcttcaaacagcctggtcg
atttcgggctgagttggagaaagtgaagctaatactgaagaggaaaggtcatgagtaatcttgcaacagtt
acaccgataaaacctcatctggaggttgtggagcatcgcgtggcagaactcgacgatggctacacccggac
tgcaaatacactgctggaagctgtcatgctttctgggcttactcaacatcagctactgattgttatggctg
tgtggcgcaagacatacggttataacaaaaaaatagattggatcggaaatgaacagttcgctgaactcact
ggcatggcgccaaccaaatgttctaccgccaaaaacgagcttatcagaatgggggttctcactcaggtggg
gcgtcaggttggtatgaataaaaatatttccgagtggaagacgaaggttaacggattcggtaaaacattta
ccagatcggtaaaactaaccttcaccaaatcggtaaaaaccaatttaccgaatcagtcaaacacaaaagac
aatatacaaaagacaataaatacaaatacccccttacccccctaaaggggggatgcgatgaaggttctaaacc
tgaaaagcgaaaacctaccaagattaactacagcgaatatcttgctgcctacaacgagattgttggtgaca
gactcccacatgcagtggaggtcaattctgaacgacaacgcaagttgaaaaagctgattgattcactggca
accaaaaacatcgacggattccgggcatacgtcaaagcgttcatggcagcagccagaccattccatttcgg
tgataacgaccgtgactgggtagctaattttgattatctgctacgcccgaaagtactgatagcaattcgtg
agggaacactatgagacaggatatcgaggcgagcgttatcggtggcttgctgattggcggattaacaccaa
ccgccagtgacgttctggcaacactggagcctgaagcattctcaattccgctctaccggaaagcttttgaa
gttattcgaaagcaggccagaaacaggaacctgattgatggactgatggtggccgaggagtgcggggatga
atacgcaacggcggtgatgatgactgcgcggtcatgtcccagcgctgcaaacctgaaaggttatgccggaa
tggttgcagacagttatcaacggcgtcaggttttacagctactggatgagatgcgggagccaatcagtaac
ggcacgctggacgcatcaggcagagcgatggacgagcttgtaaagcgcctgtcatccatcaggaagccgcg
gaacgaggttaaacctgtgcgactgggtgaaatcatcaatgactacactgacacgcttgacaggcgtctga
ggaacggagaagagtcggataccctgaagaccggaatcgaagagcttgacgctatcaccggagggatgaac
gcagaagaccttgtgattattgctgctcgtccaggtatgggtaaaaccgaactggcgctgaagatagccga
aggcgtggcaagtcgtgttattcctggttctggcgtccggcgcggtgtgttgattttctcgatggaaatga
gcgccattcaggttgttgagagagggattgccggcgcaggaatgatgtcggtcagtgtgctgcgtaacccg
tcacgtatggacgatgaaggatgggcgagagttgcaagcgggatgaagttgctggcagagctggatgtgtg
ggtagttgacgcatcgcgtttgtctgtcgaagaaatcaggtccatttccgaacgccacaagcaggagcatc
ctaatctgtcactgattatggctgactatctcgggctaattgagaaaccaaaagcggaacgtaatgacctc
gccatagcacatatctccggtagcctgaaagcgatggcgaaagacctgaaaactccagttatctccctaag
ccagctctcccgcgatgttgagaagcggccaaacaagcgcccgacaaacgcagatttgcgggattcaggaa
gcattgaacaggacgcagactcaatcatcatgctctatcgggaagcggtatatgacgagaacagtagcgcc
gcgccatttgctgaaatcatcgtgacgaaaaaccgttttggctcgcttggtacggtttaccagcggttctg
caacggacactttgttgcatgtgaccaggacgaagccagacagatttgcacggcatcaaatgcacctgctg
gacgcagaaagcgatatgcacaaggggctgacgtatgactatttacatcactgagttggtaacaggcctgc
tggtaatcgcaggcctttttatttggggggagagggaagtcatgaaaaaactaacctttgaaattcgatctc
cagcacatcagcaaaacgctattcacgcagtacagcaaatccttccagacccaaccaaaccaatcgtagta
accattcaggaacgcaaccgcagcttagaccaaaacaggaagctatgggcctgcttaggtgacgtctctcg
tcaggttgaatggcatggtcgctggctggatgcagaaagctggaagtgtgtgtttaccgcagcattaaagc
agcaggatgttgttcctaaccttgccgggaatggctttgtggtaataggccagtcaaccagcaggatgcgt
gtaggcgaatttgcggagctattagagcttatacaggcattcggtacagagcgtggcgttaagtggtcaga
cgaagcgagactggctctggagtggaaagcgagatggggagacagggctgcatga
